# Partitioned coalescence support reveals biases in species-tree methods and detects gene trees that determine phylogenomic conflicts

**DOI:** 10.1101/461699

**Authors:** John Gatesy, Daniel B. Sloan, Jessica M. Warren, Richard H. Baker, Mark P. Simmons, Mark S. Springer

## Abstract

Genomic datasets sometimes support unconventional or conflicting phylogenetic relationships when different tree-building methods are applied. Coherent interpretations of such results are enabled by partitioning support for controversial relationships among the constituent genes of a phylogenomic dataset. For the supermatrix (= concatenation) approach, several simple methods that measure the distribution of support and conflict among loci were introduced over 15 years ago. More recently, partitioned coalescence support (PCS) was developed for phylogenetic coalescence methods that account for incomplete lineage sorting and use the summed fits of gene trees to estimate the species tree. Here, we automate computation of PCS to permit application of this index to genome-scale matrices that include hundreds of loci. Reanalyses of four phylogenomic datasets for amniotes, land plants, skinks, and angiosperms demonstrate how PCS scores can be used to: 1) compare conflicting results favored by alternative coalescence methods, 2) identify outlier gene trees that have a disproportionate influence on the resolution of contentious relationships, 3) assess the effects of missing data in species-trees analysis, and 4) clarify biases in commonly-implemented coalescence methods and support indices. We show that key phylogenomic conclusions from these analyses often hinge on just a few gene trees and that results can be driven by specific biases of a particular coalescence method and/or the extreme weight placed on gene trees with high taxon sampling. Attributing exceptionally high weight to some gene trees and very low weight to other gene trees counters the basic logic of phylogenomic coalescence analysis; even clades in species trees with high support according to commonly used indices (likelihood-ratio test, bootstrap, Bayesian local posterior probability) can be unstable to the removal of only one or two gene trees with high PCS. Computer simulations cannot adequately describe all of the contingencies and complexities of empirical genetic data. PCS scores complement simulation work by providing specific insights into a particular dataset given the assumptions of the phylogenetic coalescence method that is applied. In combination with standard measures of nodal support, PCS provides a more complete understanding of the overall genomic evidence for contested evolutionary relationships in species trees.

## 1. Introduction

A hope is that phylogenomic approaches will resolve longstanding systematic controversies, in particular, rapid radiations among divergent taxa that are deep in the Tree of Life (e.g., Jarvis et al., 2014). When the relative support for different hypotheses is nearly equal, however, even genome-scale data will not solve the most challenging phylogenetic problems without attention to detail, application of the most appropriate methods, and rigorous analysis. A small bias of a given systematic method can be amplified as more data are added, especially when sequence divergences are great (e.g., Jeffroy et al., 2006; Simmons and Gatesy, 2015). Furthermore, at tightly spaced internodes, even a few homology errors (unrecognized paralogy, mis-annotation of genes, local misalignment) or a small subset of problematic outlier genes can undermine the wealth of information in large phylogenetic matrices (e.g., Brown and Thomson, 2017). The overwhelming size of modern datasets can actually hide damaging problems that would have been detected easily in the past when much smaller datasets were the norm (Philippe et al., 2017).

By partitioning support for a particular clade to the various genes in a phylogenomic dataset, the positive versus negative evidence for competing relationships can be quantified and visualized, and particularly influential loci that determine phylogenetic results can be discerned. In the supermatrix (concatenation) approach to systematics (Miyamoto, 1985; Kluge, 1989; Nixon and Carpenter, 1996; de Queiroz and Gatesy, 2007), simple methods were developed over a decade ago to characterize the distribution of supporting evidence for a particular clade. These methodologies include partitioned branch support (Baker and DeSalle, 1997), nodal dataset influence (Gatesy et al., 1999a), dataset removal index (Gatesy et al., 1999a), partitioned likelihood support (Lee and Hugall, 2003), and partition addition bootstrap alteration (Struck et al., 2006). Taxonomic congruence approaches (Nelson, 1979; Miyamoto and Fitch, 1985) that assess clade support via agreements and conflicts with independently estimated gene trees also have been used to assess corroboration for relationships in supermatrix trees (Rokas et al., 2003; Salichos and Rokas, 2013; Salichos et al., 2014; Smith et al., 2015; Kobert et al., 2016; Arcila et al., 2017).

With the recognition that bootstrapping provides an incomplete summary of support for phylogenomic datasets (Bayzid et al., 2015; Sayyari and Mirarab, 2016; Gatesy et al., 2017; Simmons et al., accepted), recent analyses of hundreds of genes have renewed interest in the partitioning of support for a particular clade among genes in genome-scale supermatrices. Shen et al. (2017) generalized partitioned likelihood support (Lee and Hugall, 2003), a method that was first applied to genome-scale supermatrices by Gatesy and Baker (2005), to comparisons between any two competing topological alternatives, and Brown and Thomson (2017) recently isolated conflicting support among loci for particular clades in a Bayesian supermatrix context. It might be expected that just a few aberrant outlier genes or homology errors in genome-scale datasets would ‘come out in the wash’ simply because a large quantity of ‘good’ data should overwhelm a few problematic genes in such large supermatrices. Empirical studies that applied partitioned support measures have demonstrated that this expectation is not necessarily the case. A partitioning of support can identify particularly influential genes that drive phylogenetic results (Gatesy et al., 1999a; Springer and Gatesy, 2018a), and the removal of just a few genes from analysis can rearrange relationships supported by supermatrices that include hundreds of genes (Brown and Thomson, 2017; Shen et al., 2017).

In addition to the supermatrix approach, more recently developed ‘summary’ coalescence methods are computationally tractable for the analysis of genome-scale systematic datasets (reviewed in Edwards, 2009; Liu et al., 2009a, 2015a). For some of the most commonly used methods, gene trees are decomposed into partially redundant counts of rooted triplets (MP-EST; Liu et al., 2010) or unrooted quartets (ASTRAL; Mirarab et al., 2014). The summed fits of gene trees to different species trees determine the optimality score (pseudo-likelihood or number of compatible quartets) and the choice among competing topologies. In such methods, the support for a given clade can be divided among the constituent gene trees for the dataset via ‘partitioned coalescence support’ (PCS) to quantify the impact of each gene tree for a particular summary coalescence method (Gatesy et al., 2017). PCS is analogous to partitioned branch support (Baker and DeSalle, 1997) and partitioned likelihood support (Lee and Hugall, 2003) in supermatrix analyses. For a mammalian dataset of 26 genes and 169 taxa, Gatesy et al. (2017) demonstrated that the relative contribution of a given gene tree to support for a particular clade can differ radically when alternative summary coalescence methods are applied and that PCS can identify problematic outlier gene trees (Fig. 1), but did not automate calculation of PCS scores for genome-scale datasets.

**Figure 1.**
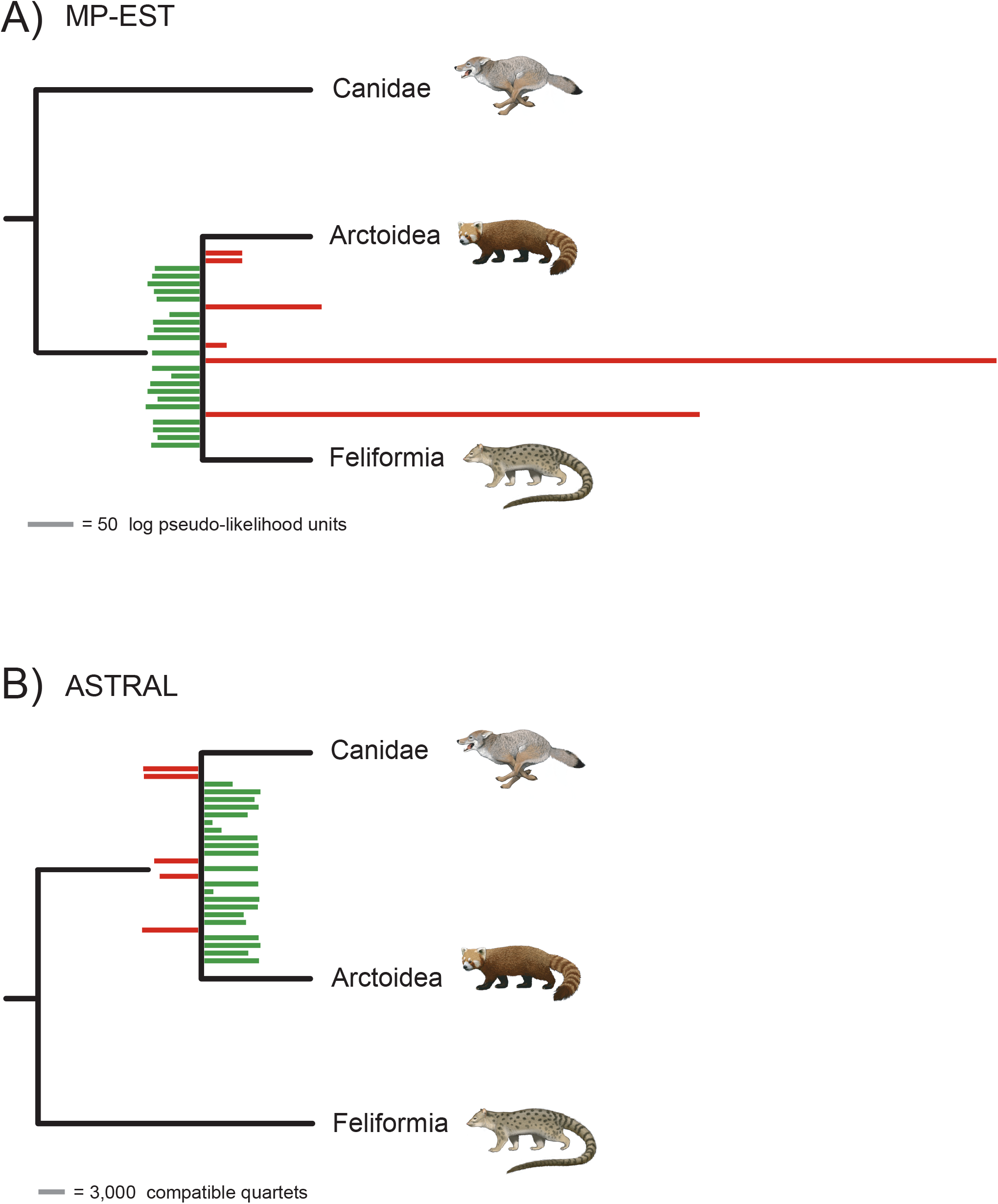
Partitioned-coalescence-support (PCS) scores reveal biased MP-EST support for Feliformia + Arctoidea (A) in contrast to coherent ASTRAL PCS support (B) for the traditional clade Caniformia (Canidae + Arctoidea). MP-EST and ASTRAL PCS scores for 26 gene trees are shown for conflicting relationships among carnivoran mammals, with positive PCS scores to the right and negative PCS scores to the left (Gatesy et al., 2017). Red bars indicate preference for the controversial Feliformia + Arctoidea resolution, and green bars indicate preference for the traditional Caniformia hypothesis. Most gene trees (21 for ASTRAL, 20 for MP-EST) support monophyly of Caniformia for both summary coalescence methods, but there are two extremely high MP-EST PCS scores for the controversial Feliformia + Arctoidea clade. When these two outlier gene trees (*EDG1*, *RAG1*) are removed from MP-EST analysis, Caniformia is supported as in the ASTRAL species tree (Gatesy et al., 2017). Magnitudes of positive and negative PCS scores are indicated in log pseudo-likelihood units (MP-EST) or compatible quartets (ASTRAL). Species trees are reduced to show the conflict between methods, but the full species trees include 169 taxa (Meredith et al., 2011).

Recent phylogenomic work increasingly has featured summary coalescence analyses of hundreds to thousands of gene trees in hopes of finally resolving the most vexing phylogenetic puzzles where incomplete lineage sorting (ILS) might hinder a supermatrix approach. As for analyses of large supermatrices (Gatesy and Baker, 2005; Brown and Thomson, 2017; Shen et al., 2017), application of partitioned support indices in genome-scale coalescence studies might enable important insights regarding conflicting support for contentious relationships. Here, we automate calculation of PCS (Gatesy et al., 2017) for genome-scale datasets and generalize the approach to comparisons between any two alternative species trees. For four published phylogenomic datasets, we use PCS to examine the distribution of support among hundreds of gene trees at controversial nodes to demonstrate the overall utility of the approach for dissecting complex signals in genomic datasets. Additionally, we utilize PCS output to compare alternative summary coalescence methods in cases where contrasting relationships are supported and where one of the coalescence methods agrees with supermatrix analysis of the same phylogenomic data. We summarize common patterns observed across the four datasets and argue for utilization of support metrics that are logically consistent with the phylogenomic coalescence methods that are applied.

## 2. Materials and Methods

### 2.1 Generalization of PCS to comparisons between any two trees and ‘scaled PCS’

Partitioned coalescence support can be calculated for summary coalescence methods, such as ASTRAL and MP-EST, that choose a species tree based on the summed fits of gene trees to different species trees (Gatesy et al., 2017). The statistic quantifies the distribution of support among gene trees for a particular clade. PCS can be positive, negative, or zero, indicating support, conflict, or ambiguity, respectively. For a fully resolved species tree, the collapse of any single internal branch results in a trichotomy, and there are three possible bifurcating resolutions at each trichotomy. Given that one of these resolutions is supported in the optimal species tree (i.e., the tree with the highest pseudo-likelihood or best quartet score), the question then becomes, how many gene trees fit this optimal species tree better than either of the two suboptimal bifurcating resolutions at this trichotomy? A PCS score for a supported clade represents comparisons between the optimal species tree that includes that clade with the two suboptimal species trees that are a single local branch swap from the optimal tree and do not support the clade. PCS for any given gene tree is the difference between the fit of that gene tree to the optimal species tree and the fit of the same gene tree to the two suboptimal species-tree topologies. For each gene tree, the alternative suboptimal species tree with best fit to the gene tree is utilized in calculating PCS, and the magnitude of the PCS score is proportional to the degree of support or conflict relative to other gene trees in the analysis (Gatesy et al., 2017). PCS scores are expressed as differences in log pseudo-likelihood scores between alternative species trees (MP-EST) or differences in the number of unrooted quartets in gene trees that fit alternative species trees (ASTRAL).

PCS scores, as described above, are determined by comparing the fits of gene trees to the optimal species tree and two suboptimal alternative topologies, but partitioned support measures can be generalized to comparisons between the optimal species tree and any conflicting species tree topology (Gatesy et al., 1999a; Shen et al., 2017). For partitioned branch support and partitioned likelihood support, comparisons are generally made between two sets of trees: the optimal supermatrix tree(s) that includes the clade of interest and the best scoring suboptimal supermatrix tree(s) that lacks the clade of interest (Baker and DeSalle, 1997, Lee and Hugall, 2003). The same could be done for PCS in a coalescence context (e.g., using ASTRAL v. 5.6.2). Alternatively, for a large dataset, a phylogenetic analysis might contradict a traditionally-recognized clade, and a comparison of among-gene support between the optimal tree(s) that lacks the traditional clade and the best suboptimal tree(s) that includes the clade might be of interest (e.g., Gatesy et al., 1999a; their figure 16). Because PCS scores summarize differences in the fits of gene trees to competing species trees, it is straightforward to make such comparisons of fit between a gene tree and any two species trees to examine differences in the distribution of support for incongruent clades.

When just two species tree topologies are compared, a PCS score also can be expressed as a *proportion* of the total difference in fit between the two species trees. For MP-EST and ASTRAL, the sum of all positive and negative PCS scores for a clade is equal to the total difference in fit between two species-tree topologies (= ‘coalescence support’ [CS] of Gatesy et al., 2017). PCS for a gene tree divided by CS, which we term ‘scaled PCS,’ therefore measures the impact that a particular gene tree has in determining the total difference in fit between two species trees. For example, if CS is 500 quartets for a comparison of two ASTRAL species-tree topologies, and PCS for a given gene tree is +250, then scaled PCS for this gene tree would be +0.5. Scaled PCS also can be greater than 1.0, zero, negative, or less than −1.0. For example, if CS is 500 quartets, and PCS for a gene tree is +750 quartets, then scaled PCS is +1.5. For this same example, if PCS for a gene tree is 0 then scaled PCS is 0; if PCS is −100 then scaled PCS is −0.2; and if PCS is −1000 then scaled PCS is −2.0. For a particular clade, the sum of scaled PCS scores across all genes in a dataset will equal 1. A scaled PCS score of >1 for a gene tree therefore indicates that the influence of a single gene rivals the entire sum of support for a clade. Henceforth we use ‘standard’ PCS scores in most analyses in this paper. For plots that show the influence of gene tree size in MP-EST versus ASTRAL analyses, we use scaled PCS (see section 3.2.2 *Gene trees with extensive missing taxa are severely downweighted in coalescence analyses).*

### 2.2 Published datasets that support conflicting phylogenomic resolutions

We investigated the utility of PCS by examining four phylogenomic datasets that include taxa with deep divergences in the Tree of Life (Fig. 2; Chiari et al., 2012; Zhong et al., 2013; Xi et al., 2014; Linkem et al., 2016). MP-EST and ASTRAL reanalyses of optimal gene trees (i.e., gene trees based on the original sequence alignments—not bootstrapped data) were performed in each case to derive the optimal species tree (i.e., the species tree with the highest optimality score for the coalescence method that was applied). We then calculated PCS scores for conflicting clades that were central to the primary phylogenetic inference in each case study by comparing the fits of gene trees to the optimal species tree and to an alternative suboptimal species tree. A brief summary of each dataset is given here, including number of taxa, outgroups, number of genes, key conflicting clades, and the species trees used in PCS calculations. More detailed context for each empirical example is given in the Results and Discussion section.

**Figure 2.**
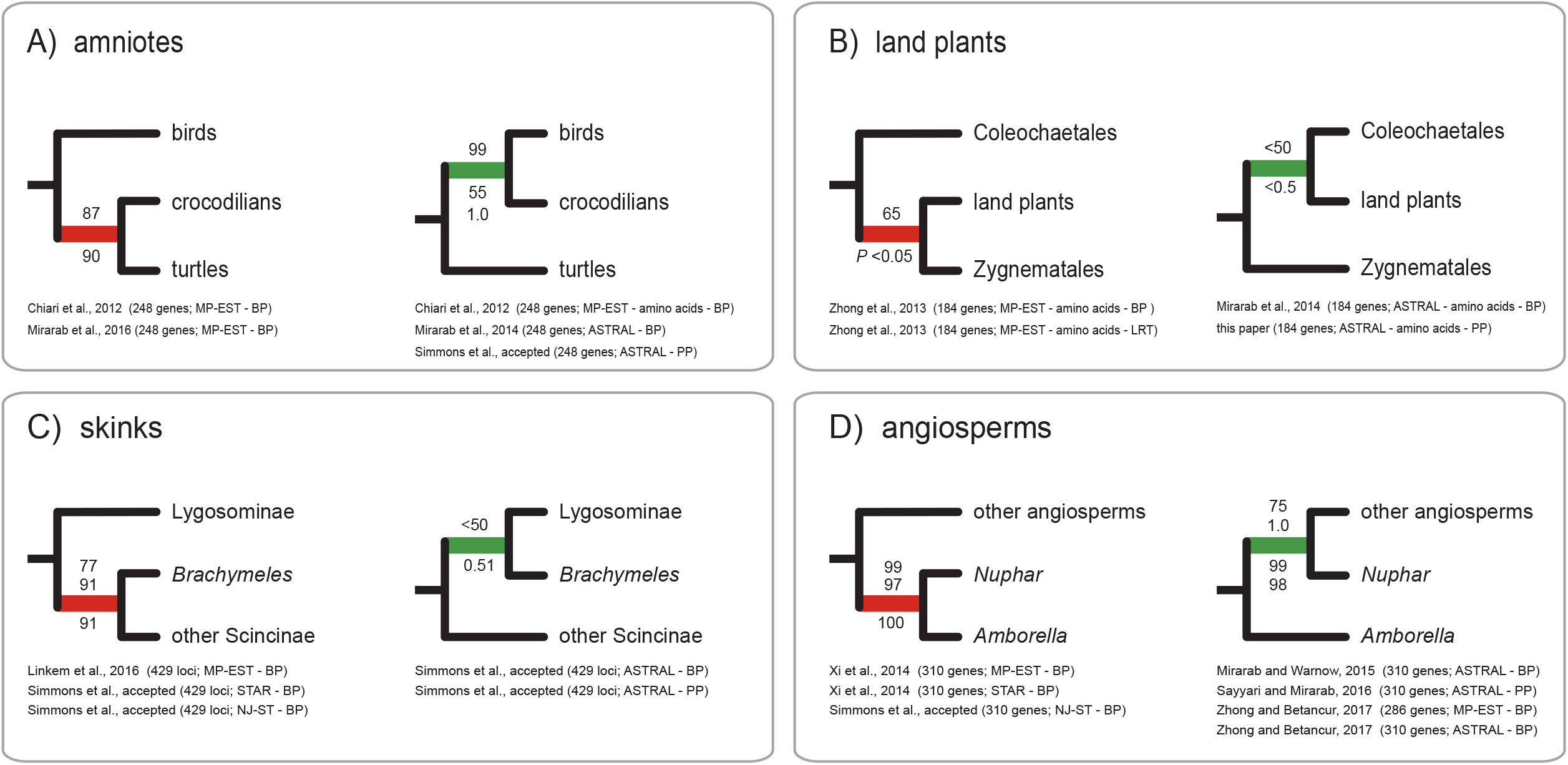
Conflicting relationships at controversial nodes for the four phylogenomic datasets that were reanalyzed here: A) amniotes (Chiari et al., 2012), B) land plants (Zhong et al., 2013), C) skinks (Linkem et al., 2016), and D) angiosperms (Xi et al., 2014). For each dataset, alternative resolutions are supported by MP-EST and ASTRAL. Support scores are shown at internodes (likelihood ratio test = LRT, bootstrap = BP, Bayesian local posterior probability = PP). For the angiosperms dataset, MP-EST analysis of optimal gene trees supports *Amborella* sister to *Nuphar* + other angiosperms, but bootstrap support is 99% for the conflicting *Amborella* + *Nuphar* clade

1. <u>Amniote Phylogeny</u> (Chiari et al., 2012): 248 protein-coding genes, 14 ingroup taxa (birds, crocodilians, turtles, lizards, mammals), and two outgroups (frog, fish). For optimal gene trees based on DNA sequences (Chiari et al., 2012), PCS scores were calculated for ASTRAL and MP-EST species trees at conflicting resolutions among crocodilians, turtles, and birds. For each coalescence method, two topologies were contrasted: crocodilians sister to birds and crocodilians sister to turtles (Fig. 2A).
2. <u>Land-Plant Origins</u> (Zhong et al., 2013): 184 protein-coding genes, 20 genera of land plants, nine genera of streptophyte algae, and an outgroup clade (three chlorophytes). For optimal gene trees based on amino-acid sequences (Zhong et al., 2013), PCS scores were calculated for ASTRAL and MP-EST species trees for conflicting resolutions among land plants, Zygnematales, and Coleochaetales. For each coalescence method, two topologies were contrasted: Zygnematales sister to land plants and Coleochaetales sister to land plants (Fig. 2B).
3. <u>‘Anomaly Zone’ in Skink Phylogeny</u> (Linkem et al., 2016): 429 ultraconserved element (UCE) loci, 15 ingroup taxa (scincid lizards), and a single outgroup (*Xantusia* lizard); every gene was sampled for each taxon. For optimal gene trees based on DNA sequences (Linkem et al., 2016), PCS scores were calculated for ASTRAL and MP-EST species trees at conflicting resolutions among Lygosominae, the scincine *Brachymeles, and a* clade composed of the other six scincine species in the dataset. For each coalescence method, two topologies were contrasted: *Brachymeles* sister to other members of Scincinae and *Brachymeles sister to Lygosominae (Fig. 2C).*
4. <u>Basal-Angiosperm Relationships</u> (Xi et al., 2014): 310 protein-coding genes, 42 ingroup taxa (angiosperms), and four outgroup taxa (three gymnosperms and *Selaginella*). For optimal gene trees based on DNA sequences (Xi et al., 2014), PCS scores were calculated for ASTRAL and MP-EST species trees at conflicting resolutions among the angiosperm *Amborella*, the angiosperm *Nuphar*, and a clade composed of the other 40 angiosperms in the dataset. For each coalescence method, two topologies were contrasted: *Amborella* sister to other angiosperms and the clade of *Amborella* + *Nuphar* sister to the remaining angiosperms (Fig. 2D).

### 2.3 Automation of partitioned coalescence support (PCS) for genome-scale datasets

For PCS calculations, an optimal species trees and one or more alternative suboptimal species trees are required input. Optimal species trees were estimated using ASTRAL 4.11.1 (Mirarab and Warnow, 2015) and MP-EST 1.5 (Liu et al., 2010). For ASTRAL 4.11.1 runs, analyses were performed with default settings using optimal gene trees from each of the four empirical studies (Chiari et al., 2012; Zhong et al., 2013; Xi et al., 2014; Linkem et al., 2016) to estimate the optimal species tree for each dataset. The same four sets of gene trees were employed in analyses using MP-EST 1.5 with default search parameters. Gene trees were rooted as in the original studies. For each dataset, 1000 independent searches for the optimal MP-EST species tree were done. For optimal species trees derived from three of the four empirical datasets, MP-EST and ASTRAL trees conflicted at the key nodes summarized in Figure 2A-C. For the fourth dataset (Xi et al., 2014), the optimal MP-EST and ASTRAL trees agree in supporting *Amborella* sister to other angiosperms, but bootstrap support in MP-EST analysis is 99% for a conflicting species tree in which *Amborella* is sister to *Nuphar* (Fig. 2D; Xi et al., 2014). The *Amborella* + *Nuphar clade also received high bootstrap support in STAR (97%) and* NJ-ST (100%) coalescence analyses (Fig. 2D). Xi et al. (2014) interpreted this as the correct topology, therefore this resolution was chosen as the alternative relationship for PCS calculations.

For each optimal species tree that was estimated, the suboptimal species-tree resolution was constructed by manually making a single local branch swap in the optimal species tree. For the optimal species tree and the alternative suboptimal species tree, branch-length information was then removed. ASTRAL species trees in this format were used as input for PCS calculations because branch lengths do not impact the optimality score for ASTRAL (the fit of quartets in gene trees to alternative species tree topologies). Because branch lengths (in coalescent units) are integral to the optimality criterion of MP-EST (pseudo-likelihood scores based on fit of rooted triplets to alternative species tree topologies), branch lengths for optimal and suboptimal MP-EST species trees were optimized using the “2” command of MP-EST 1.5 by running at least three independent searches for each species tree (see Gatesy et al., 2017).

For PCS calculations at conflicting nodes, the optimal and suboptimal species trees described above were used as input along with optimal gene trees from the empirical studies. PCS calculations were as in Gatesy et al. (2017) and were automated with custom Perl scripts (pcs_mpest.pl and pcs_astral.pl). Theses scripts take a set of gene trees as input, and the user must also specify the optimal species tree and at least one alternative topology. The scripts iterate over the set of gene trees, calling the respective program to calculate log pseudo-likelihood scores (MP-EST) or quartet scores (ASTRAL) based on comparisons between the gene tree and each of the optimal/alternative species-tree topologies. The scripts then parse the resulting output and calculate PCS scores as defined above. For each gene tree, they report the best-supported species tree and the calculated scores. For each of the four published datasets, PCS scores were calculated for conflicting MP-EST and ASTRAL species trees using these scripts. Code for automating PCS analysis is freely available in the pcs_mpest and pcs_astral repositories at https://github.com/dbsloan.

Bootstrapping (Felsenstein, 1985) and Bayesian local posterior probabilities (PPs) (Sayyari and Mirarab, 2016) were used to assess support for species-tree relationships in comparisons to PCS. Bootstrap support for species trees was taken from published studies of the four datasets that were reanalyzed here. ASTRAL was used to calculate Bayesian local PP, a measure of clade support that does not entail resampling of nucleotide sequence data and is instead based on analysis of optimal gene trees (Sayyari and Mirarab, 2016). ASTRAL 4.11.1 calculates Bayesian local PP for each clade in the optimal ASTRAL species tree. For alternative clades that are not resolved in the best species tree, ASTRAL 5.6.1 (Zhang et al., 2017) was used to infer Bayesian local PPs.

## 3. Results and Discussion

### 3.1 Examples of PCS for four phylogenomic datasets with conflicting resolutions

Here, we present our results as four separate vignettes that demonstrate the utility of PCS scores for understanding complex phylogenetic signals that emerge in summary coalescence analyses of genome-scale datasets that include hundreds of genes. Supplementary data and results (MP-EST control files, gene trees, species trees, species trees with alternative resolutions at critical nodes, and PCS scores for each dataset) are posted at: https://figshare.com/s/b0a7cdbc7f3c63737920. This is currently a private link; it will be changed to a permanent public posting).

#### 3.1.1 A few gene trees undermine robust support for genome-scale species trees of Amniota

Chiari et al. (2012) analyzed a phylogenomic dataset comprised of 248 genes from 14 amniote taxa (birds, crocodilians, turtles, lizards, mammals), a close outgroup (the frog *Xenopus*), and a more distant outgroup (the fish *Protopterus*). They alternatively analyzed amino-acid as well as DNA sequences using both coalescence and concatenation approaches. Most, but not all, concatenation analyses robustly support the traditional clade Archosauria (crocodilians + birds) to the exclusion of turtles. MP-EST coalescence analyses of gene trees based on the amino-acid sequences corroborate this result with high bootstrap support (99%), but MP-EST analysis of gene trees based on DNA sequences group turtles and crocodilians as sister taxa with 87%-90% bootstrap support (Chiari et al., 2012; Mirarab et al., 2016). This DNA-based result represents a large swing in MP-EST bootstrap support for this controversial phylogenetic hypothesis (Fig. 2A). Subsequent ASTRAL coalescence reanalyses of the DNA sequences contradict the MPEST coalescence result and favor monophyly of Archosauria with 55% bootstrap support (Mirarab et al., 2014) and Bayesian local PP of 1.0 (Simmons et al., accepted). The relatively strong MP-EST nucleotide bootstrap support for the incongruent result (crocodilians + turtles) shows a shift in topology relative to ASTRAL coalescence analysis that yields maximum PP for Archosauria (Fig. 2A). For gene trees based on DNA sequences, we calculated PCS scores for ASTRAL and MP-EST species trees at the conflicting nodes. In each case, we compared the two discrepant results: Archosauria vs. crocodilians sister to turtles.

Given the relatively high MP-EST bootstrap score for the crocodilians + turtles clade (Fig. 2A), the difference between pseudo-likelihood scores for the two contrasting phylogenetic hypotheses is surprisingly small, 7.949 log likelihood units (Fig. 3). For MP-EST, there are 112 positive PCS scores (mean positive PCS score = +2.205 log likelihood units) for the crocodilians + turtles clade that is supported by this coalescence method, while there are 136 negative PCS scores (mean negative PCS score = −1.757 log likelihood units) for Archosauria. That is, within the context of MP-EST analysis, 24 more gene trees favor the traditional Archosauria clade than the controversial crocodilians + turtles clade (Fig. 3A). Likewise, more gene trees (79 of 248) support Archosauria than the crocodilians + turtles clade (60 of 248).

**Figure 3.**
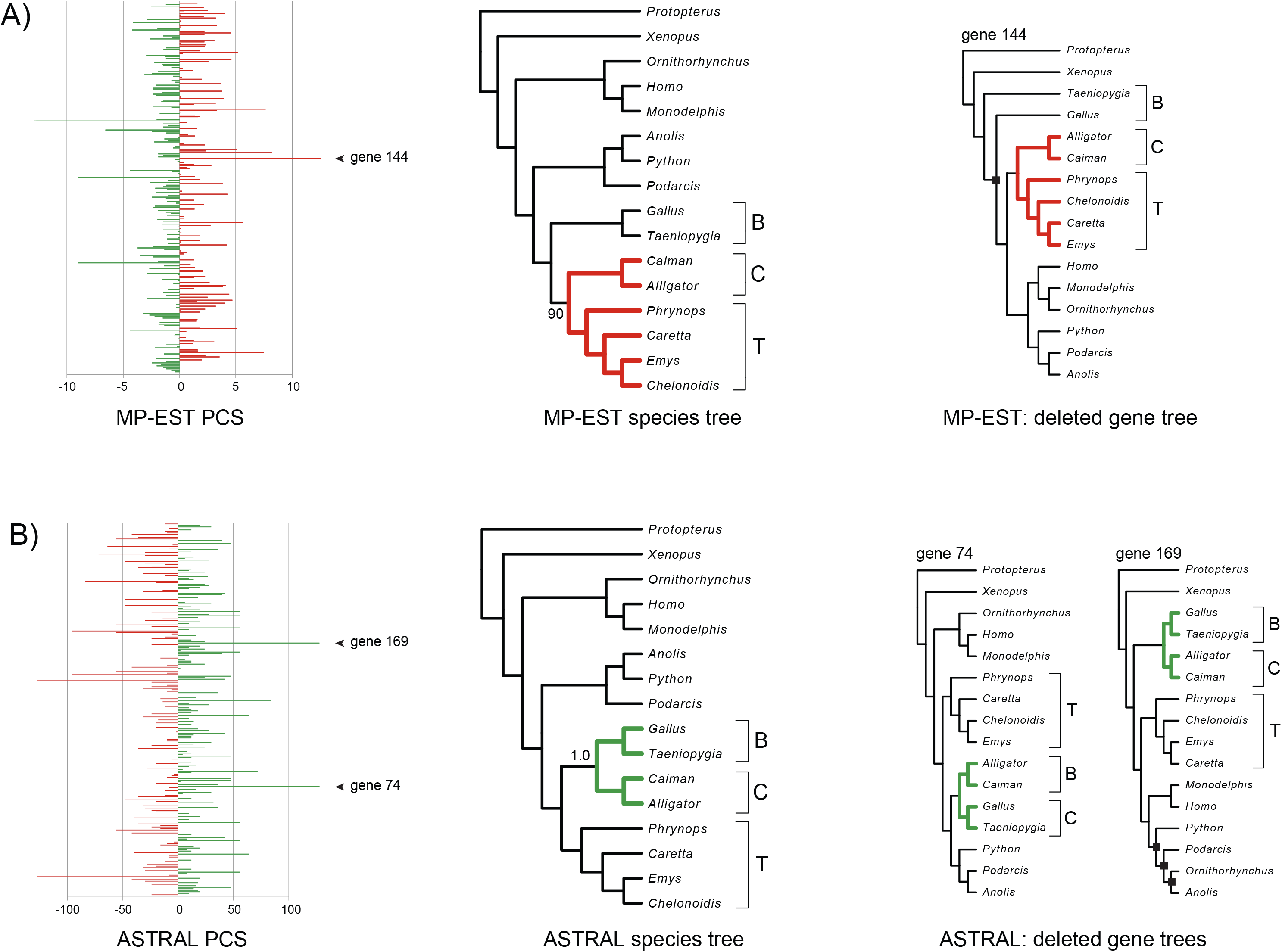
MP-EST (A) and ASTRAL (B) partitioned coalescence support (PCS) scores and instability to removal of high-PCS gene trees for the amniotes dataset (Chiari et al., 2012). For both MP-EST and ASTRAL, the following are shown from left to right: PCS scores for all gene trees, optimal species tree based on all gene trees, and high-PCS gene trees whose removal resulted in loss of the focal clade (red = crocodilians + turtles; green = crocodilians + birds [Archosauria]). Bootstrap support (MP-EST) and Bayesian local posterior probability (ASTRAL) are shown for relationships among crocodilians, birds, and turtles. The black square in gene tree #144 marks the clade that conflicts with monophyly of birds; in gene tree #169, black squares mark clades that conflict with both Squamata (lizards and snakes) and Mammalia (mammals). Abbreviations are: C = crocodilians, B = birds, T = turtles.

The range of MP-EST PCS scores is wide, and several outlier gene trees with extremely high or extremely low PCS scores are apparent. For MP-EST analysis, gene tree #144 has the highest positive PCS score of +12.525 log likelihood units for the crocodilians + turtles clade. Gene tree #144 represents an implausible topology in which Aves (birds) is paraphyletic due to a misrooting of the amniote tree on *Taeniopygia* (finch) (Fig. 3A). Given recent estimates of divergence times (Meredith et al., 2011), gene tree #144 would require retention of ancestral polymorphism for >200 MY, which is not credible unless extreme balancing selection at this locus was sustained over this long time frame (Piertney and Oliver, 2006; Leffler et al., 2013). This gene tree does, however, support the crocodilians + turtles clade. After removal of this gene tree from the dataset, MP-EST reanalysis of the remaining 247 gene trees produces a species tree that contradicts the original MP-EST analysis and instead supports the traditional clade Archosauria. Thus, support for the controversial crocodilians + turtles clade hinges on just a single gene tree that was identified by PCS, and the distribution of support among gene trees suggests that Archosauria is better supported.

In a previous reanalysis of the DNA sequence data from Chiari et al. (2012), Brown and Thomson (2015) executed Bayesian supermatrix analyses of the same 248 genes and partitioned support for conflicting resolutions among crocodilians, birds, and turtles using Bayes factors. Their analyses revealed two clear outlier genes, #30 and #59, which were both impacted by paralogy problems and were highly influential in their Bayesian supermatrix analysis. Gene trees derived from these two loci support the controversial crocodilians + turtles clade that is favored by MP-EST analysis. For both genes, MP-EST PCS scores are not extremely high (gene #30 = +5.106, gene #59 = +4.091), but the sum of these two PCS scores, +9.197 log likelihood units, exceeds the difference in log likelihoods between the crocodilians + turtles tree and the Archosauria tree. After removal of these two gene trees from the amniote dataset, MP-EST reanalysis of the remaining 246 optimal gene trees again supports a monophyletic Archosauria in contrast to the MP-EST analysis of all 248 gene trees. Thus, the crocodilians + turtles result for MP-EST is also sensitive to the removal of these two problematic gene trees.

For ASTRAL analysis of optimal gene trees from Chiari et al. (2012), phylogenetic results are more compatible with PCS scores and congruence among gene trees. There are 124 positive ASTRAL PCS scores (mean positive PCS score = +24.750 quartets) for Archosauria, and there are 104 negative PCS scores (mean negative PCS score = −28.105 quartets) for the crocodilians + turtles clade (Fig. 3B). Twenty gene trees show no preference for either of these two hypotheses (i.e., PCS = 0). Archosauria is favored by ASTRAL analysis with a local PP of 1.0 (Fig. 2A), indicating maximum Bayesian support for this traditional clade, but the difference in quartet scores for the two alternative hypotheses (Archosauria versus crocodilians + turtles) is relatively low (146 quartets) and ASTRAL bootstrap support also is low (55%; Fig. 2A). As for MP-EST, there are several gene trees with very high or very low PCS scores. Gene tree #74 (PCS = +128 quartets) and gene tree #169 (PCS = +128 quartets) have the highest ASTRAL PCS at this node, and each score approaches the difference between quartet scores for the two competing phylogenetic hypotheses (+146 quartets). When we removed gene tree #74 and reanalyzed the remaining 247 gene trees in ASTRAL, Archosauria is still supported with a local PP of 1.0. Likewise, when we removed just gene tree #169 and reanalyzed the remaining 247 gene trees in ASTRAL, Archosauria is again supported with a local PP of 1.0. However, when we removed both of these gene trees, ASTRAL analysis of the remaining 246 gene trees instead supports a species tree in which the crocodilians + turtles is resolved but Bayesian local PP is 0.0 for this clade (Fig. 3B). Hence, deletion of the second gene tree from ASTRAL analysis rearranges the topology and shifts support from a local PP of 1.0 for the supported Archosauria clade (247 gene trees) to a local PP of 0.0 for the supported crocodilians + turtles clade (246 gene trees), a dramatic change that hinges on a single locus.

By partitioning support among gene trees, PCS scores can be used to identify influential gene trees that tip support for one phylogenetic hypothesis over another in summary coalescence analyses (Fig. 3), but PCS scores additionally show how missing taxa in different gene trees can impact interpretations of support in MP-EST and ASTRAL species trees. Both methods work by decomposing gene trees down to smaller units, rooted triplets for MP-EST and unrooted quartets for ASTRAL, and then utilizing this information to reconstruct the species tree based on the fit of these small subtrees to competing species trees hypotheses. It is widely acknowledged that missing taxa in gene trees can impact phylogenetic results when using these and other summary coalescence methods (Liu et al., 2010; Hovmöller et al., 2013; Mirarab et al., 2014; Springer and Gatesy, 2014; Zhong et al., 2014; Hosner et al., 2016; Xi and Davis, 2016). A serious problem is that gene trees which include many taxa contain more partially redundant triplets or quartets than do gene trees with fewer taxa.

Because PCS measures differences in gene-tree support for alternative hypotheses within the context of the optimality criterion for the coalescence method that is applied, the specific effects of different gene tree sizes (with respect to numbers of taxa sampled) can be clearly quantified. Figure 4 shows two gene trees that both support monophyly of Archosauria but with very different numbers of sampled taxa. Gene tree #169 includes all 16 taxa that were sampled by Chiari et al. (2012) and is one of the two gene trees with the highest ASTRAL PCS score (+128) for Archosauria. By contrast, gene tree #56 is characterized by a much lower PCS score (+10) and includes only nine of 16 taxa (Fig. 4). Despite no conflicts with the ASTRAL species tree, gene tree #56 has a PCS score that is 12.8 times lower than the PCS score for gene tree #169 at the Archosauria node. This difference is caused by the many more unrooted quartets in gene tree #169 relative to gene tree #56 that are informative regarding relationships among crocodilians, birds, and turtles (Fig. 4). Thus, in an ASTRAL analysis, a large gene tree like #169 will have more influence than 12 small gene trees like #56. For MP-EST, the difference between PCS scores is likewise skewed for the two gene trees in this example. Because the crocodilians + turtles clade is supported by MP-EST analysis, however, the MP-EST PCS scores are negative instead of positive: −12.897 for the big gene tree #169 and just −0.991 for the small gene tree #56 (Figs. 3, 4). So for MP-EST, gene tree #169 has >13 times more negative influence at the crocodilians + turtles node relative to the much smaller gene tree #56.

**Figure 4.**
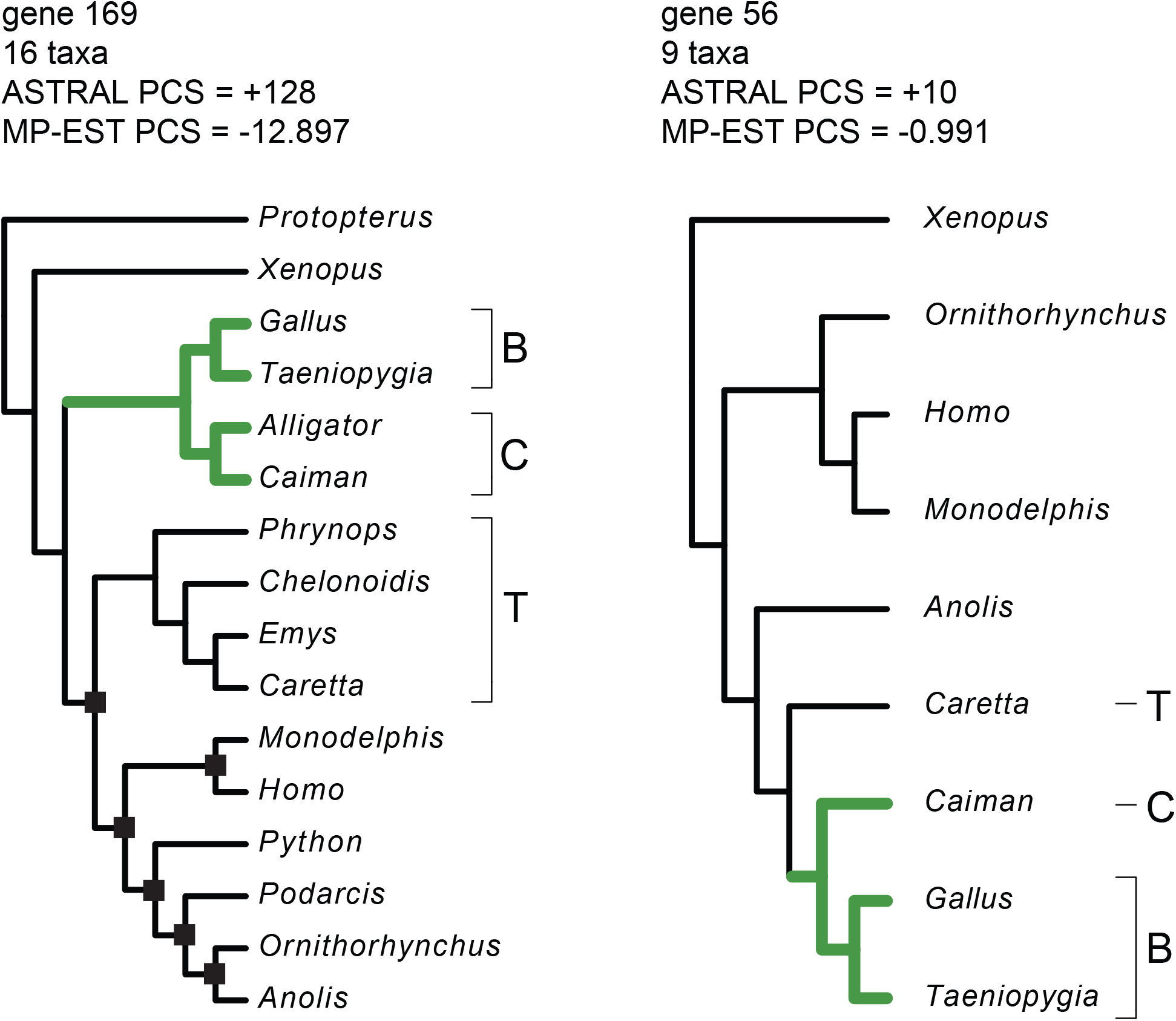
Disparities in partitioned-coalescence-support (PCS) scores for two gene trees with different numbers of taxa for the amniotes dataset (Chiari et al., 2012). PCS scores for both ASTRAL and MP-EST are shown. Black squares indicate nodes that conflict with the ASTRAL species tree (Fig. 3B). Note that according to PCS scores, the large gene tree (#169) has >13 times more influence relative to the small gene tree (#56) for the comparison between conflicting resolutions (Fig. 2A). The crocodilians + birds clade (Archosauria) is marked by green branches. Abbreviations are: C = crocodilians, B = birds, T = turtles.

Given that in the coalescence approach to systematics, each independently-sorting locus is treated as the basic unit of analysis (Doyle, 1997; Maddison, 1997; Slowinski and Page, 1999; Edwards, 2009), it is problematic that different gene trees can differ in their impact by an order of magnitude simply because more species are sampled for one gene than for another. In the present case, genes like #144 that support the crocodiles + turtles resolution (Fig. 3A) are outnumbered by genes like #56 (Fig. 4) that support Archosauria according to both PCS and taxonomic congruence (i.e., counts of gene trees that support each group), but PCS scores for genes that favor the crocodilians + turtles clade are higher, on average, in both MP-EST and ASTRAL analyses (Fig. 3). We contend that there is no reason to overweight the outlier gene tree #144 to such an extent simply because more taxa were sampled for this gene. PCS scores can help quantify the effects of missing data in any case study by providing dataset-specific information on the magnitude of supporting and conflicting evidence in gene trees for the summary coalescence method that is applied (Figs. 3, 4).

Missing taxa also impact interpretations of support in the ASTRAL analysis due to a discrepancy between the optimality criterion for ASTRAL and the method used to quantify nodal support. ASTRAL analysis of all 248 gene trees supports the traditional clade Archosauria. More gene trees support this resolution relative to the crocodilians + turtles clade according to PCS, and Bayesian local PP for Archosauria is high (1.0). This result is easily overturned, however, by removing just two out of 248 gene trees from the ASTRAL analysis (Fig. 3B). When Bayesian support is assessed for the crocodilians + turtles clade that is favored by the analysis of 246 gene trees (i.e., two gene trees removed), the local PP of 0.0 indicates *no* Bayesian support for a hypothesis that is preferred by the ASTRAL analysis. The “-t 2” option of ASTRAL 5.6.1 (Zhang et al., 2017) permits more precise calculation of Bayesian local PPs for all three alternative resolutions at a trichotomy. ASTRAL 5.6.1 outputs 0.9965 local PP for Archosauria and just 0.0034 local PP for the crocodilians + turtles clade that is supported by the analysis of 246 gene trees.

While ASTRAL simply tabulates the *numbers of gene-tree quartets* that fit alternative species tree hypotheses to choose a best tree, the Bayesian local PP method instead treats *each informative gene tree* as the basic unit of analysis and compares three possible resolutions of a trichotomy (Sayyari and Mirarab, 2016). Because more gene trees actually support Archosauria relative to the crocodilians + turtles clade, even when two gene trees that support Archosauria are removed and archosaur monophyly is lost in the ASTRAL species tree, it is possible to have a high Bayesian local PP of 0.9965 for Archosauria. The criterion for choosing the optimal ASTRAL species tree is different from the criterion for measuring support using Bayesian local PP. Therefore, paradoxical situations like the one described above can emerge when conflicts and missing taxa in a phylogenomic dataset are extensive. ASTRAL bootstrap analysis avoids this inconsistency between how a tree is chosen and how support for that tree is measured. However, ASTRAL (like MP-EST) is prone to unjustified weighting of different gene trees by an order of magnitude or more when taxa are incompletely sampled (Figs. 3B, 4), and this weighting by gene tree size also applies to ASTRAL bootstrap analysis (but not Bayesian local PP).

#### 3.1.2 Phylogenomic coalescence analyses of land plant origins are impacted by missing taxa

In a phylogenomic investigation of land plant origins, Zhong et al. (2013) analyzed a dataset of 184 loci from 20 land-plant genera, representatives from several clades of streptophyte algae (9 genera), and an outgroup clade (three chlorophyte genera). MP-EST coalescence analysis of this dataset supports a sister-group relationship between land plants and Zygnematales algae with moderate support (65% bootstrap). MP-EST bootstrap analyses of smaller and larger samplings of genes (42, 78, 119, 211, or 289 genes) with various amounts of missing taxa uniformly support this same relationship (bootstrap up to 61%; Fig. 2B). Zhong et al. (2013) suggested that the consistency of MP-EST results and the more erratic results for concatenation imply that the MP-EST coalescence tree is accurate. Although MP-EST bootstrap support was low, no higher than 65%, MP-EST likelihood ratio tests for the 184-gene dataset indicate solid support for Zygnematales sister to land plants and rejection of three viable alternative hypotheses at *P* < 0.05 (Zhong et al., 2013, 2014). Reanalyses of the same gene trees (Springer and Gatesy, 2014) showed that STAR coalescence bootstrap analyses instead support Zygnematales + Coleochaetales as the algae sister group to land plants for five of the six datasets (bootstrap up to 82%), a hypothesis that agrees with ML concatenation bootstrap trees for two of the six datasets (Zhong et al., 2013). Mirarab al. al. (2014) reanalyzed the preferred 184 locus dataset using ASTRAL, and this produced a third species tree, Coleochaetales algae sister to land plants, which is weakly supported (bootstrap <50%; Fig. 2B). We compared PCS scores for alternative species trees by comparing the MP-EST coalescence resolution in which Zygnematales is sister to land plants to the ASTRAL coalescence resolution in which Coleochaetales are sister to land plants (Fig. 2B).

MP-EST coalescence analysis of 184 optimal gene trees (Fig. 5A) supports a species tree that positions Zygnematales algae as sister to land plants as in the published MP-EST bootstrap analysis (Zhong et al., 2013). The difference between the log pseudo-likelihood for this hypothesis and the alternative resolution in which Coleochaetales are sister to land plants is 73.320 log likelihood units. The number of MP-EST PCS scores for the Coleochaetales-sister-toland-plants resolution (102 negative PCS scores) is greater than for the preferred Zygnematales-sister-to-land-plants hypothesis (82 positive PCS scores). Given the high degree of gene-tree conflict in this dataset (Simmons et al., 2016), the great majority of independently-estimated gene trees support neither hypothesis. Seventeen gene trees resolve Zygnematales sister to land plants, and just 13 gene trees resolve Coleochaetales sister to land plants.

**Figure 5.**
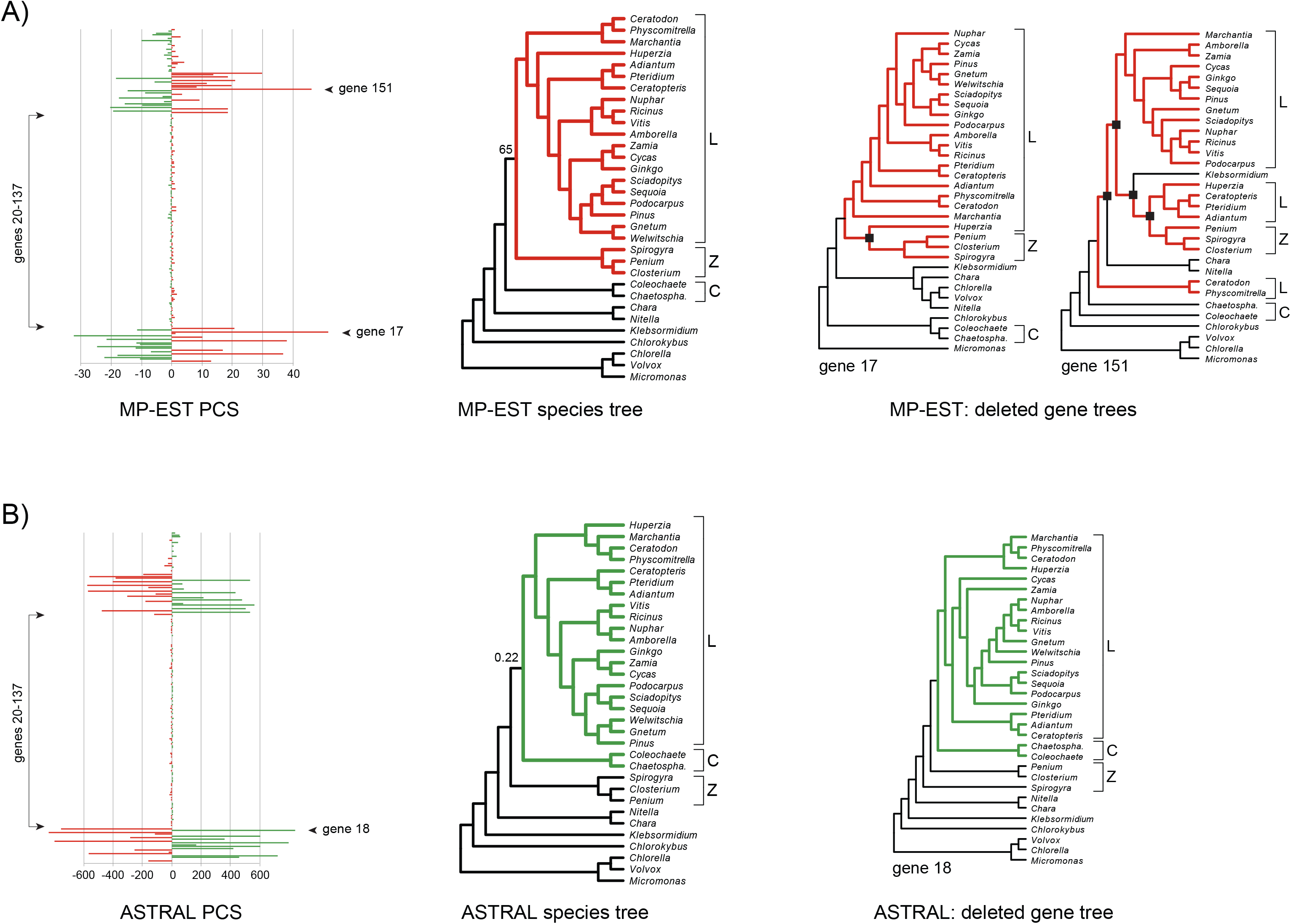
MP-EST (A) and ASTRAL (B) partitioned-coalescence-support (PCS) scores and instability to removal of high-PCS gene trees for the land-plants dataset (Zhong et al., 2013). For both MP-EST and ASTRAL, the following are shown from left to right: PCS scores for all gene trees, optimal species tree based on all gene trees, and high-PCS gene trees that were removed to effect loss of the focal clade (red = land plants + Zygnematales; green = land plants + Coleochaetales). For the two gene trees with highest MP-EST PCS, various algae lineages are nested within paraphyletic land plants (black squares at nodes). Note that small gene trees which include <8 taxa (genes #20-137) have extremely low PCS scores for both MP-EST and ASTRAL analyses. The sum of ASTRAL PCS scores (absolute values) for the 118 small gene trees is just 461 quartets (average = 3.907 quartets), which is much less than ASTRAL PCS for the single large gene tree #18 (+840 quartets). Bootstrap support for the land plants + Zygnematales clade and Bayesian local posterior probability for the land plants + Coleochaetales clade are shown. Abbreviations are: L = land plants, Z = Zygnematales, C = Coleochaetales, *Chaetospha*. = *Chaetosphaeridum*.

For MP-EST, PCS scores vary widely with most genes trees characterized by very low scores (absolute values) and a few gene trees with high PCS in the positive or negative direction (Fig. 5A). To a large extent, these differences are caused by extensive missing taxa in the gene trees. Missing taxa are not randomly distributed in the land-plants dataset. The same 24 taxa are absent from 65% of the gene trees (#s 20-137). The 118 small gene trees (< 8 taxa) provide low PCS support in MP-EST analysis relative to the larger gene trees in the dataset (Fig. 5A). For example, the large gene tree #12, which includes 31 taxa, supports Zygnematales as sister to land plants, and has the third highest MP-EST PCS score (+36.916) for this relationship. By contrast, the small gene tree #38, which includes just 8 taxa, also supports Zygnematales sister to land plants, but has a low PCS score (+1.624). Because there are few rooted triplets in the small gene tree, the much larger gene tree has nearly 23 times more influence in the MP-EST analysis.

In the original description of the MP-EST method, Liu et al. (2010; pp. 4-5) noted that, missing data “in some gene trees are allowed if lineages are missing randomly, but a lot of missing lineages may dramatically reduce the performance of the pseudo-likelihood approach.” Zhong et al. (2014; p. 270) later defended the utility of MP-EST for coalescence analysis of the land-plants dataset despite the extent and uneven distribution of missing data, asserting that, “We observed a considerable number of missing taxa in the plant data… Thus we chose the MP-EST method [Liu et al., 2010] rather than the STAR method, to reduce the effect of missing data on species tree estimation, because MP-EST is based on gene tree triples, which are comparable among all gene trees.” Given so few rooted triplets in the 118 small gene trees, however, these gene trees cannot exert much influence. PCS scores for the small gene trees (#s 20-137) are barely visible in the overall plot of MP-EST PCS (Fig. 5A).

According to PCS, two outlier gene trees have high impacts in the MP-EST analysis (Fig. 5A). The large gene tree #17 includes all 32 taxa in the dataset and has the highest positive PCS score (+50.292) for the preferred MP-EST resolution (Zygnematales sister to land plants), but for this gene tree, Zygnematales groups within land plants, rendering the latter paraphyletic. The large gene tree #151 also includes all 32 taxa and has the next highest PCS score (+44.738), but Zygnematales and two additional algae lineages, *Klebsormidium* and Charales (*Chara*, *Nitella*), are nested within land-plants (Fig. 5A). When these two large gene trees that do not even support land-plant monophyly are removed, MP-EST analysis of the remaining 182 gene trees instead favors a species tree in which Coleochaetales are the streptophyte algae clade that is sister to land plants. This result demonstrates that the MP-EST result hinges on just two large outlier gene trees.

ASTRAL coalescence analysis of the 184 gene trees in the land-plants dataset produced a different species tree (Fig. 5B) that positions Coleochaetales algae sister to land plants (Mirarab et al., 2014). The difference in quartet scores between this hypothesis and the Zygnematales-sister to-land-plants hypothesis is 677 quartets. Bayesian local PP for the preferred resolution of Coleochaetales sister to land plants is just 0.22 (Figs. 2B, 5B), and local PP is actually higher for the Zygnematales-sister to-land-plants resolution (0.69). The distribution of ASTRAL PCS scores also indicates a preference for Zygnematales sister to land plants (76 negative PCS scores) relative to the best-scoring ASTRAL hypothesis (72 positive PCS scores), with 36 equivocal gene trees (PCS = 0; Fig. 5B).

Because of the large blocks of missing taxa, ASTRAL PCS scores have an overall pattern that is similar to MP-EST PCS scores. Small gene trees with < 8 taxa have uniformly low PCS scores, while large gene trees can have very high positive PCS or very negative PCS (Fig. 5B). The implication is that large gene trees are highly upweighted relative to small gene trees in ASTRAL coalescence analysis as well. For example, the large gene tree #18 includes all 32 taxa, supports Coleochaetales sister to land plants, and has the highest ASTRAL PCS for the Coleochaetales + land plants clade (+840; Fig. 6). By contrast, the small gene tree #107 includes just six taxa, also supports Coleochaetales + land plants, but has a tiny PCS score for this clade (+3). Because there are just 15 unrooted quartets in a small gene tree, the large gene tree with 35,960 unrooted quartets has 280 times more influence in ASTRAL coalescence analysis than the small gene tree (Fig. 6). Given that 118 of the 184 gene trees in the land plants dataset are small (< 8 taxa; #s 20-137), all of these small gene trees combined have less influence in ASTRAL analysis relative to the single large gene tree #18 at the Coleochaetales + land plants node (Fig. 5B). When the 118 small gene trees are analyzed in isolation from the larger gene trees, the ASTRAL species tree agrees with the species tree supported by MP-EST analysis of all 184 gene trees (Zygnematales sister to land plants), with 0.76 PP. By contrast, ASTRAL analysis of the 66 larger gene trees favors a conflicting resolution in which Coleochaetales is sister to land plants (0.51 local PP), which matches the ASTRAL species tree for all 184 gene trees (Figs. 2B, 5).

**Figure 6.**
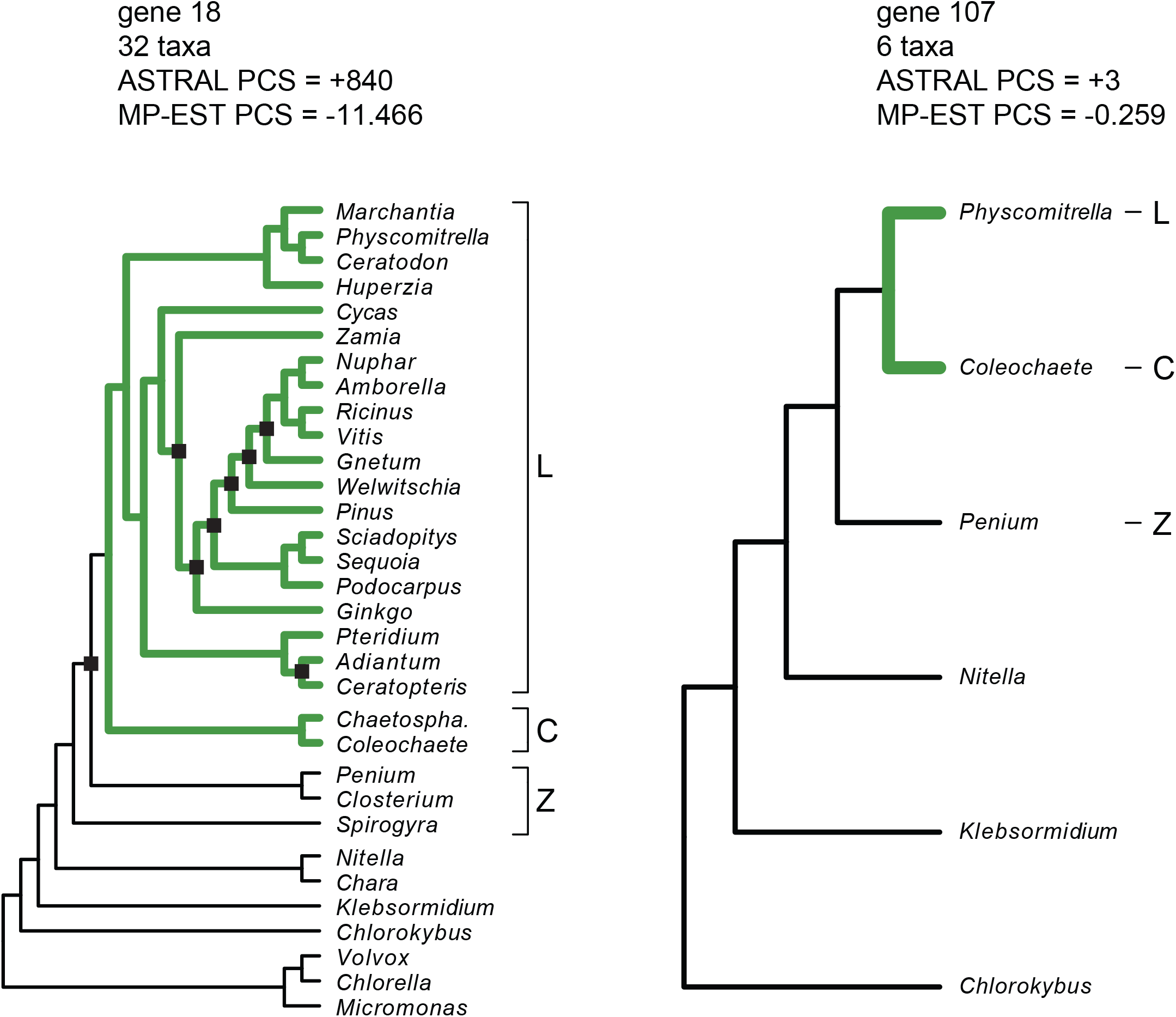
Disparities in partitioned-coalescence-support (PCS) scores for gene trees with different numbers of taxa for the land-plants dataset (Zhong et al., 2013). PCS scores for both ASTRAL and MP-EST are shown. Black squares mark nodes in gene trees that conflict with the ASTRAL species tree (Fig. 5B). Note that the large gene tree #18 (32 taxa) has 280 times more influence in ASTRAL analysis relative to the perfectly-congruent small gene tree #107 (6 taxa) and >44 times more influence in MP-EST analysis. For a gene tree with 32 taxa, there are 4,960 rooted triplets and 35,960 unrooted quartets, but for a gene tree with six taxa, there are just 20 triplets and 15 quartets. The land plants + Coleochaetales clade is indicated by green branches. Abbreviations are: L = land plants, Z = Zygnematales, C = Coleochaetales, *Chaetospha*. = *Chaetosphaeridum*.

The ASTRAL species tree is even less stable than the MP-EST species tree that is based on the same 184 gene trees. When the large gene tree #18 (PCS = +840 quartets) is removed (Fig. 5B), ASTRAL analysis of the remaining 183 gene trees favors the sister-group relationship supported by MP-EST (Zygnematales + land plants). Thus, for MP-EST and ASTRAL, support shifts back and forth between two alternative resolutions following small perturbations in gene sampling, and neither method agrees with a third coalescence method, STAR, that instead supports Zygnematales + Coleochaetales sister to land plants (Springer and Gatesy, 2014). Three different coalescence methods yield three different results for this same dataset, and bootstrap-support scores are low in all cases (Zhong et al., 2013; Mirarab et al., 2014; Springer and Gatesy, 2014). Zhong et al. (2013, 2014) argued that despite low MP-EST bootstrap support (65%) for the Zygnematales + land plants clade (Fig. 2B), the MP-EST log pseudo-likelihood score for this hypothesis versus alternatives was compelling for the 184-gene dataset based on significant likelihood-ratio tests. Here, we have found that the highly skewed taxon sampling in different gene trees created a situation where alternative phylogenetic hypotheses are supported by different coalescence methods depending on the overweighted influence of just a few large gene trees (Fig. 5). As for the amniote dataset (Figs. 3, 4), PCS scores for the land-plants dataset discern sensitivity to the removal of just a few gene trees from analysis as well as the striking effects of missing taxa in summary coalescence analyses (Figs. 5, 6).

#### 3.1.3 The ‘anomaly zone’ and conflicting phylogenomic analyses of skinks

Linkem et al. (2016) analyzed a phylogenomic dataset of 429 ultraconserved element (UCE) loci from 15 scincid lizards and a single outgroup (*Xantusia*); all 429 genes were sampled for each taxon. The mean conflict between gene trees in the skinks dataset is remarkably high, an average of 11.2 of 13 clades disagree in pairwise comparisons (Simmons et al., accepted). Coalescence (MP-EST) and concatenation results disagree where there are several short internodes in phylogenetic reconstructions. Linkem et al. (2016) hypothesized that the MP-EST coalescence tree, which supports Scincinae (77% bootstrap; Fig. 2C), is accurate and that the maximum likelihood (ML) supermatrix analysis that strongly contradicts this clade and instead favors *Brachymeles + Lygosominae (100% bootstrap) is inaccurate. They suggested that this was* an empirical example of the ‘anomaly zone,’ a region of tree space where the most probable gene tree does not match the species tree (Degnan and Rosenberg, 2006; Kubatko and Degnan, 2007). In simulations of the ‘anomaly zone,’ concatenation can fail while summary coalescence methods can perform well when large numbers of genes are sampled (Kubatko and Degnan, 2007; but see Mendes and Hahn, 2018). Subsequent ASTRAL coalescence analysis of optimal gene trees from Linkem et al. (2016) supported the ML concatenation result (local PP = 0.51, bootstrap = <50%, but STAR (91% bootstrap for Scincinae) and NJ-ST (91% bootstrap for Scincinae) coalescence analyses agreed with MP-EST results (Simmons et al., accepted). We compared PCS scores for the alternative species trees by comparing the MP-EST coalescence resolution in which Scincinae is monophyletic to the ASTRAL coalescence resolution in which the scincine *Brachymeles* groups with Lygosominae to the exclusion of other members of Scincinae (Fig. 2C).

MP-EST coalescence analysis of 429 optimal gene trees supports monophyly of Scincinae, but PCS scores indicate that 49 more gene trees favor the contradictory sister-group relationship between *Brachymeles* and Lygosominae (239 negative PCS scores) over monophyly of Scincinae (190 positive PCS scores; Fig. 7A). The smaller proportion of positive PCS scores for Scincinae relative to the more abundant negative PCS scores for *Brachymeles* + Lygosominae corresponds to the counts of individual gene trees that resolve each of these contrasting phylogenomic hypotheses. Out of 429 gene trees, only 12 resolve monophyly of Scincinae, which is favored by MP-EST, while nearly three times as many independently estimated gene trees, 33, resolve the conflicting *Brachymeles + Lygosominae clade. Out of 429 gene trees, only* one (gene tree #174) positions *Brachymeles* sister to remaining scincines as in the optimal MPEST tree, while 21 gene trees place *Brachymeles* sister to Lygosominae, which is the preferred ML supermatrix result (Linkem et al., 2016). The difference in pseudo-likelihood scores between the optimal MP-EST species tree that supports Scincinae and the alternate species tree with the *Brachymeles + Lygosominae clade is just 14.222 log likelihood units. Given that only 12 gene* trees resolve monophyly of Scincinae while 33 resolve the conflicting *Brachymeles* + Lygosominae clade, the 384 gene trees that contradict both clades drive MP-EST’s preference for Scincinae (Fig. 7A).

**Figure 7.**
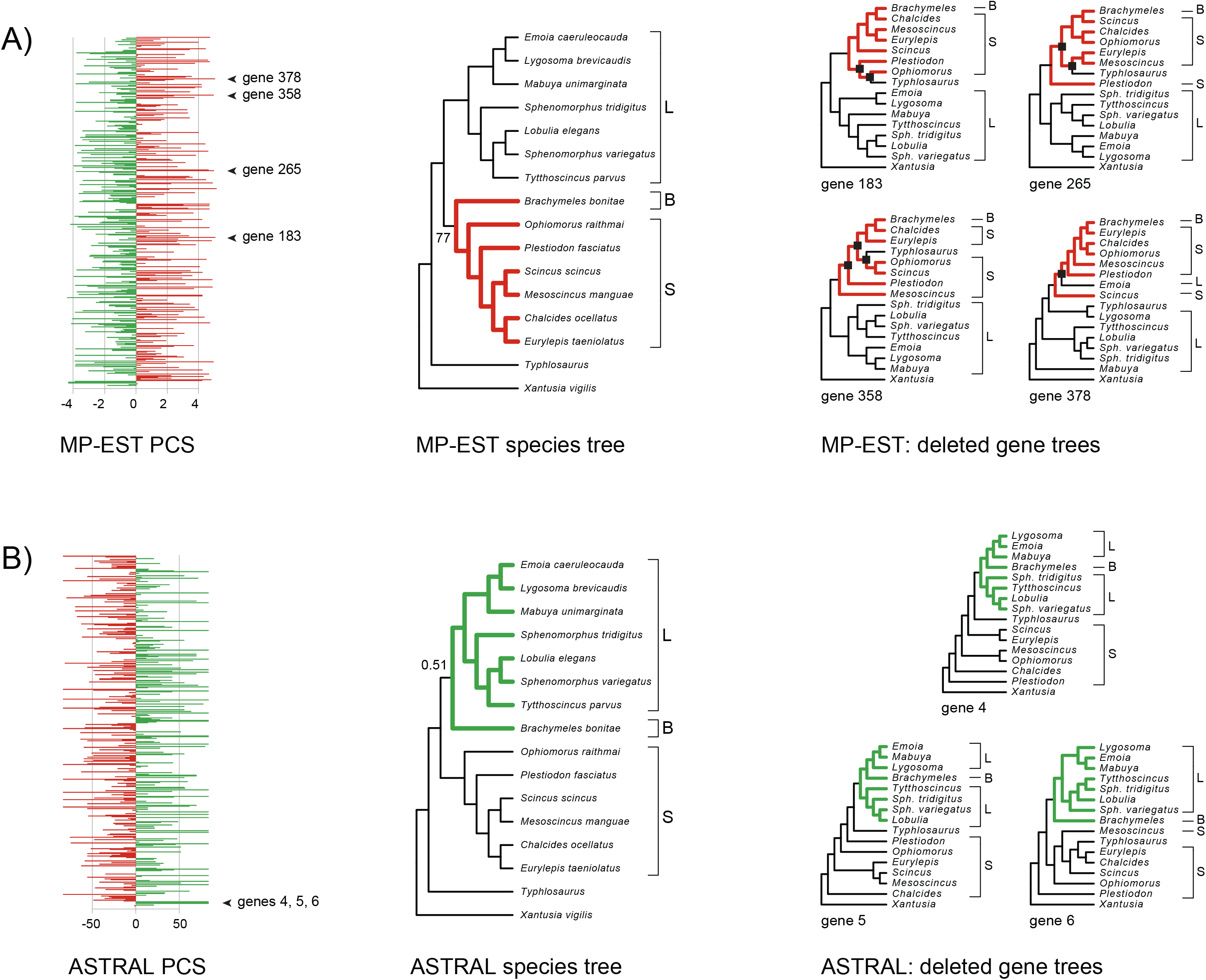
MP-EST (A) and ASTRAL (B) partitioned-coalescence-support (PCS) scores and instability to removal of high-PCS gene trees for the skinks dataset (Linkem et al., 2016). There are no missing taxa in any of the 429 gene trees for this dataset. For both MP-EST and ASTRAL, the following are shown from left to right: PCS scores for all gene trees, optimal species tree based on all gene trees, and high-PCS gene trees whose removal resulted in loss of the focal clade (red = Scincinae; green = *Brachymeles* + Lygosominae). For the four gene trees with high MPEST PCS for Scincinae that were removed, Scincinae is not even supported as a clade. Black squares mark nodes that disrupt monophyly of Scincinae in these gene trees. For the ASTRAL tree based on all 429 gene trees, Bayesian local posterior probability is shown for the *Brachymeles + Lygosominae clade, and bootstrap support for Scincinae is indicated for the MP*-EST analysis of all gene trees. Abbreviations are: L = Lygosominae, B = *Brachymeles*, S = all scincines except *Brachymeles*, *Sph*. = *Sphenomorphus.*

The ten gene trees with the highest MP-EST PCS scores for monophyly of Scincinae are shown in Figure 8. Surprisingly, seven of the ten gene trees contradict monophyly of Scincinae. *Typhlosaurus* (subfamily Acontinae) is positioned within subfamily Scincinae in six of the ten gene trees, and the critical unstable taxon, *Brachymeles, is nested well within Scincinae in all ten* of these gene trees, which is not the case for the optimal MP-EST species tree (Fig. 7A). Gene tree #174, the only one of 429 that resolves *Brachymeles* sister to other scincines as in the MPEST tree, has a lower MP-EST PCS score for Scincinae (+3.794) relative to many gene trees that contradict this clade (Fig. 8; PCS = +4.813 to +5.037). The gene trees with *Brachymeles* deeply nested among scincines have PCS scores greater than +4 (Fig. 8). We removed four of these gene trees (#183, 265, 358, 378) that also contradict a monophyletic Scincinae (Figs. 7A, 8). MP-EST analysis of the remaining 425 gene trees overturns support for Scincinae, and the alternative *Brachymeles + Lygosominae clade is supported in the resulting MP-EST tree. This species tree* is identical to the RAxML supermatrix tree presented by Linkem et al. (2016). PCS scores identify just a few gene trees, less than 1% of the total, that tip support for the contentious Scincinae clade. Surprisingly, all four gene trees that were removed contradict monophyly of Scincinae (Fig. 7A), demonstrating the instability of this clade in the preferred MP-EST tree.

**Figure 8.**
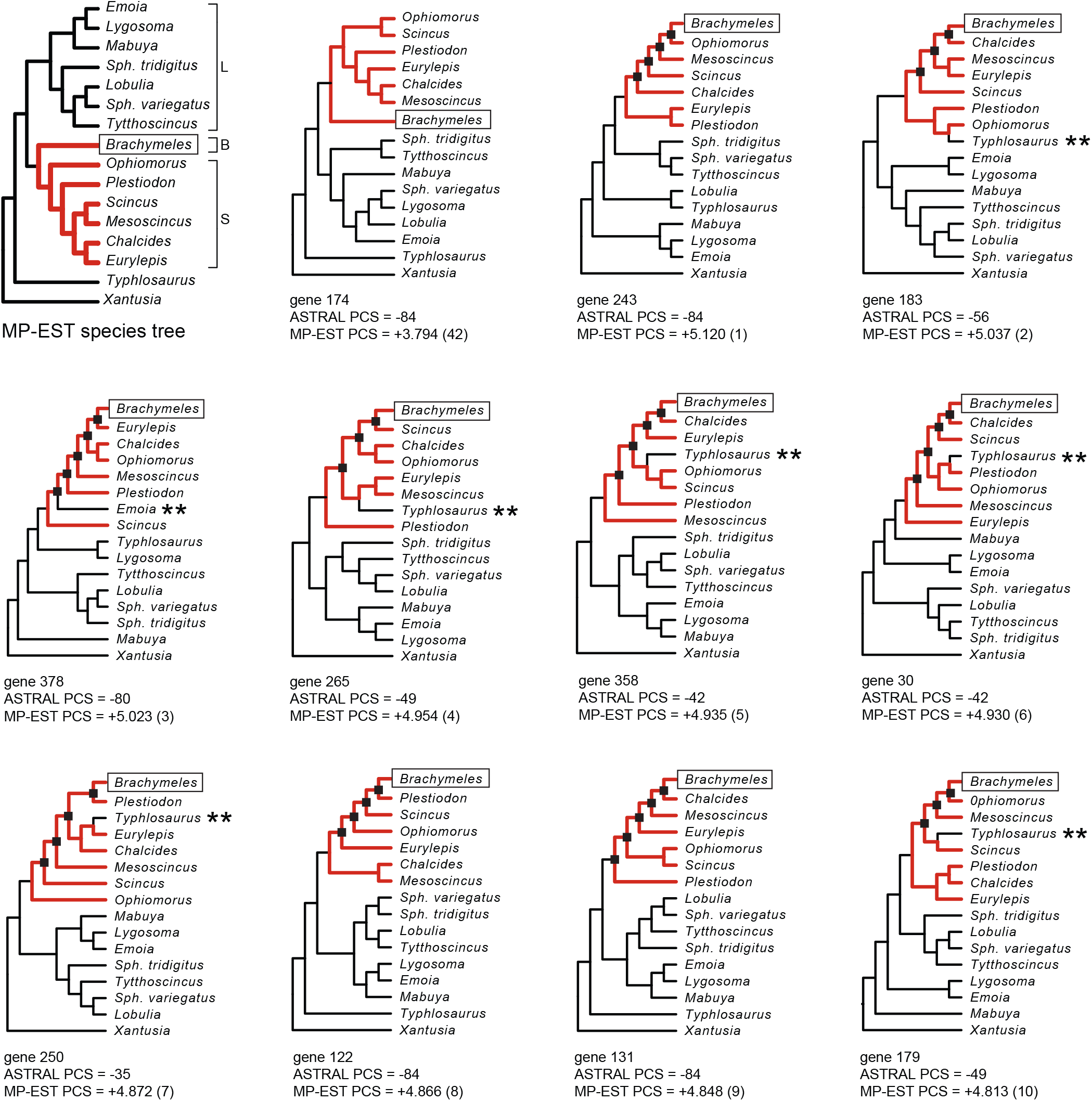
Gene trees from Linkem et al. (2016) with the top ten MP-EST PCS scores for monophyly of Scincinae. In contrast to the MP-EST species tree and a gene tree (#174) with the 42nd highest PCS score (top left), *Brachymeles is deeply nested among scincines (black squares* at nodes) in the ten gene trees with the highest MP-EST PCS. Seven of the ten gene trees contradict monophyly of Scincinae; double asterisks (**) indicate non-scincine taxa that are nested within Scincinae (red branches connect scincine taxa). The rank of each MP-EST PCS score is shown in parentheses (e.g., gene tree #183 has the second highest MP-EST PCS score). Note that gene tree #174 is the only gene tree out of 429 that positions *Brachymeles* sister to other scincines as in the MP-EST species tree. Abbreviations are: L = Lygosominae, B = *Brachymeles*, S = all scincines except *Brachymeles*, *Sph*. = *Sphenomorphus.*

In contrast to MP-EST results, ASTRAL coalescence analysis of the full set of 429 optimal gene trees supports a species tree that contradicts monophyly of Scincinae and instead favors the *Brachymeles* + Lygosominae clade that Linkem et al. (2016) suggested was an artifact of ML supermatrix analysis. In contrast to MP-EST PCS, ASTRAL PCS scores indicate that 33 more gene trees favor monophyly of Scincinae (209 negative PCS scores) than the *Brachymeles* + Lygosominae clade that is supported by ASTRAL (176 positive PCS scores), with 44 ties (PCS = 0; Fig. 7B). The distribution of ASTRAL PCS scores contrasts with the resolution of independently estimated gene trees, but the 33 gene trees that support *Brachymeles* + Lygosominae have the highest PCS scores for this clade in ASTRAL analysis. Because there are no missing taxa in gene trees, each gene tree that supports this clade has the same ASTRAL PCS score (+84 quartets). Likewise, the 12 gene trees that support Scincinae have the lowest ASTRAL PCS scores (−84; Fig. 8). The difference in overall quartet scores between the optimal ASTRAL species tree versus the alternative species tree that resolves Scincinae is 199 quartets. Removal of three gene trees with the highest PCS at this node (gene trees #4, #5, #6), and subsequent ASTRAL analysis of the remaining 426 gene trees contradicts *Brachymeles +* Lygosominae and instead supports the conflicting Scincinae clade. As for MP-EST, ASTRAL support is sensitive to the removal of just a few gene trees (Fig. 7B).

For the skinks dataset, gene trees with the highest ASTRAL PCS scores for the supported *Brachymeles + Lygosominae clade are those gene trees that resolve the Brachymeles* + Lygosominae clade (Fig. 7B). This is the expected result; clear-cut support for a particular clade in a gene tree should yield high partitioned support for that clade in phylogenetic coalescence analysis when taxon sampling is complete. For MP-EST analysis of the same 429 gene trees, however, the expectation of a match between resolution of a clade in a gene tree and high partitioned support for that gene tree in coalescence analysis does not generally hold. Instead, many of the most influential gene trees with high PCS for the supported Scincinae clade contradict a monophyletic Scincinae because *Typhlosaurus* (subfamily Acontinae) commonly groups within this clade. Furthermore, *Brachymeles*, a pivotal taxon in this dataset (Fig. 2C; Linkem et al., 2016), is placed in apical, nested positions in all of these high-PCS gene trees that sharply conflict with the optimal species tree (Fig. 8), a pattern that also emerged in MP-EST analysis of the land plants dataset (Fig. 5A). This ‘apical nesting’ of a basal lineage within its sistergroup might explain the biased upweighting of these gene trees in MP-EST analysis. The large influence of gene trees that are highly incongruent with the species tree is not a desirable attribute of the MP-EST method (also see Simmons and Gatesy, 2015; Gatesy et al., 2017).

Linkem et al. (2016: p. 468) chose to use the MP-EST coalescence method in their study “…because it can accurately estimate the species tree topology despite the anomaly zone… [Liu et al. 2010].” A complication is that the species tree based on ASTRAL coalescence analysis of these data is perfectly congruent with the ML concatenation tree but conflicts with the MP-EST coalescence tree. Thus, the two coalescence methods that explicitly account for ILS and are statistically consistent when gene trees are reconstructed accurately (Liu et al., 2010; Mirarab and Warnow, 2005) provide conflicting results (Fig. 2C). We therefore contend that the bias quantified by high MP-EST PCS scores for incongruent gene trees (Figs. 7A, 8) offers an alternative explanation for disagreements among phylogenetic methods in analyses of the skinks dataset. Given the generally slow rate of evolution in UCE loci (presumably due to negative selection) and several extremely short internal branches in the skinks species tree (Linkem et al., 2016), it is unlikely that all gene tree in this dataset have been reconstructed accurately (∼86% pairwise incongruence of gene trees). Therefore, summary coalescence methods that assume no natural selection and accurate reconstruction of gene trees (Liu et al., 2010; Mirarab et al., 2014) should not be expected to reliably resolve extremely challenging anomaly-zone conditions, especially at deep divergences in the Tree of Life (Gatesy and Springer, 2014; Springer and Gatesy, 2016). In such cases, even minor differences in how alternative coalescence methods interpret the evidence provided by incongruent gene trees (Fig. 8) can turn phylogenomic coalescence results this way (Fig. 7A) or that (Fig. 7B).

#### 3.1.4 Long-branch misrooting of angiosperm phylogeny biases coalescence results

Xi et al. (2014) analyzed a phylogenomic dataset of 310 protein-coding genes from 42 angiosperm taxa, a close outgroup clade (three gymnosperms), and a single distant outgroup (*Selaginella*). Comparisons between coalescence (MP-EST, STAR) and ML concatenation results revealed a robustly supported conflict at the base of angiosperms. The two coalescence methods group *Nuphar* with *Amborella* (MP-EST bootstrap = 99%; STAR bootstrap = 97%) at the base of angiosperms, while ML supermatrix analysis positions *Amborella* as sister to a clade of the remaining angiosperms (100% bootstrap; Fig. 2D). Xi et al. (2014) argued that the robust supermatrix result is an artifact of concatenation being misled by fast-evolving sites. Subsequently, ASTRAL coalescence reanalysis of the same dataset contradicted the *Nuphar* + *Amborella* clade and favored the supermatrix result (Simmons and Gatesy, 2015) with moderate (75% bootstrap; Mirarab and Warnow, 2015) to high (1.0 Bayesian local PP; Sayyari and Mirarab, 2016) support (Fig. 2D). Misplacement of the extremely long *Selaginella* outgroup was implicated in the conflict between coalescence methods that use rooted versus unrooted gene trees as inputs (Mirarab and Warnow, 2015; Simmons and Gatesy, 2015; Simmons, 2017). This interpretation was bolstered by coalescence analyses (MP-EST, STAR, ASTRAL) that resolve *Amborella* as sister after excluding the long-branched *Selaginella* (Simmons and Gatesy, 2015; Zhong and Betancur, 2017). For Xi et al.’s (2014) original dataset that includes 310 gene trees for 46 taxa, we compared PCS scores at the focal conflicting nodes by comparing the *Amborella* + *Nuphar* resolution to the *Amborella*-basal resolution (Fig. 2D).

Although MP-EST bootstrap support for the *Amborella* + *Nuphar* clade is 99% (Xi et al., 2014), MP-EST analysis of the 310 optimal gene trees does not even support this clade (Fig. 9A). The optimal MP-EST species tree instead supports *Amborella* sister to the remaining angiosperms, as in the supermatrix analyses (Simmons and Gatesy, 2015). This topology is just slightly better (4.368 log likelihood units) than the alternative species tree that positions *Amborella* + *Nuphar* sister to the remaining angiosperms. Bootstrap support of 99% for a clade that is contradicted by analysis of the original sequence data is a surprising result. In terms of partitioned support (Fig. 9A), there are 24 more positive PCS scores for the species tree in which *Amborella* is sister to the remaining angiosperms (167) than negative PCS scores for the species tree in which *Amborella* + *Nuphar* is sister to other angiosperms (143). This pattern agrees with independently estimated gene trees; 82 optimal gene trees support *Amborella* sister to the remaining angiosperms, but only 28 gene trees support *Amborella* + *Nuphar* as sister (Simmons et al., accepted).

**Figure 9.**
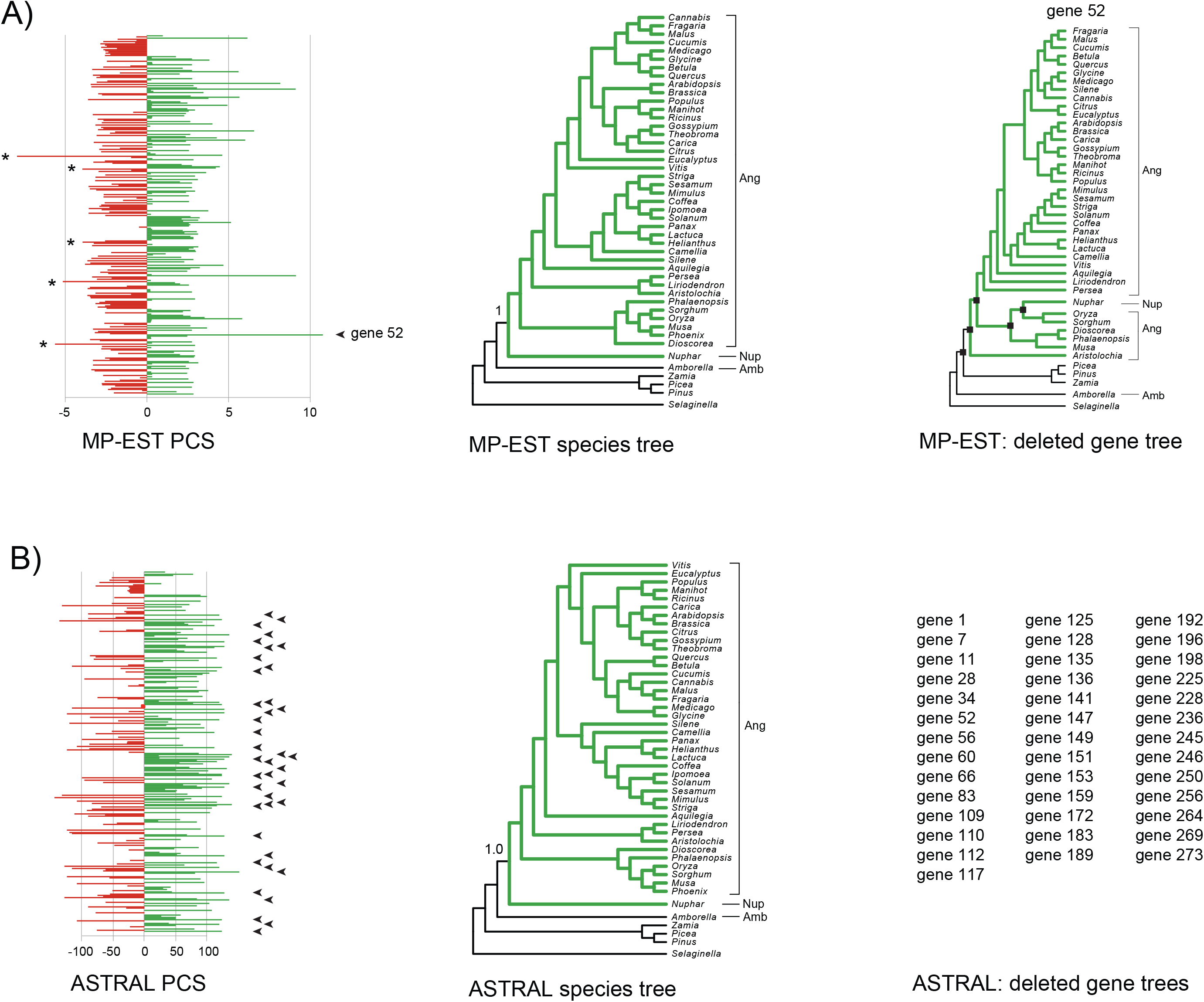
MP-EST (A) and ASTRAL (B) partitioned-coalescence-support (PCS) scores and instability to removal of high-PCS gene trees for the angiosperms dataset (Xi et al., 2014). For both MP-EST and ASTRAL, the following are shown from left to right: PCS scores for all gene trees, optimal species tree based on all gene trees, and high PCS gene trees whose removal resulted in loss of the clade that includes all angiosperms except *Amborella* (green lineages). For ASTRAL, the Bayesian local posterior probability is 1.0 for the clade composed of all angiosperms except *Amborella*, and removal of the 40 gene trees with highest PCS scores collapses this clade (only gene #s shown; arrowheads point to PCS scores for these genes). Bootstrap support is just 1% for the same clade in MP-EST analysis, and removal of a single gene tree (#52) shifts support to the conflicting *Amborella* + *Nuphar* clade. Note that *Nuphar* is nested well within angiosperms (black squares at nodes) and that angiosperms are paraphyletic for this gene tree. Single asterisks (*) identify the five gene trees with the most MP-EST support for the *Amborella* + *Nuphar* clade (red). All of these gene trees contradict the *Amborella* + *Nuphar clade* (Fig. 10). Abbreviations are: Amb = *Amborella*, Nup = *Nuphar*, Ang = all angiosperms except *Amborella* and *Nuphar.*

The distribution of MP-EST PCS scores reveals several outlier gene trees in both the negative and positive directions (Fig. 9A). Gene tree #52 has the highest MP-EST PCS score for the clade that includes all angiosperms except *Amborella* (+10.760 log likelihood units). This PCS score is more than twice the log pseudo-likelihood difference that separates the two conflicting species-tree resolutions, and is 4 times greater than the average value (+2.305) for the 167 positive MP-EST PCS scores. Gene tree #52 does support the clade of all angiosperms except *Amborella*, but gymnosperms are nested within angiosperms due to the very basal position of *Amborella* in the gene tree (Fig. 9A). This topology implies a deep coalescence of >100 MY according to molecular clock estimates (Bell et al., 2010; Magallón et al., 2013) and is more credibly interpreted as an inaccurately reconstructed gene tree (Simmons and Gatesy, 2015). The very high MP-EST PCS score for this gene tree represents another example of the ‘basal dragdown bias’ characterized previously in MP-EST but not ASTRAL analyses (Gatesy et al., 2017). Removal of this single outlier gene tree from the dataset and subsequent MP-EST analysis of the remaining 309 gene trees shifts support to a species tree that positions *Amborella* + *Nuphar* sister to remaining angiosperms (Fig. 9A). In MP-EST analyses of optimal gene trees, support is tenuous for contentious relationships at the base of angiosperms given sensitivity to the removal of a single gene tree. This contrasts with the MP-EST bootstrap consensus (Fig. 2D) that indicates robust support (99%) for the *Amborella* + *Nuphar clade.*

ASTRAL coalescence analysis of the full set of 310 optimal gene trees supports the *Amborella*-basal species tree, but in comparison to MP-EST, bootstrap support is much higher for this resolution (75% for ASTRAL versus 1% for MP-EST), and the Bayesian local PP is 1.0. The *Amborella* basal tree is 4930 quartets better than the *Amborella* + *Nuphar* basal tree according to ASTRAL, and more ASTRAL PCS scores support the former (141) over the latter (100) hypothesis, with 69 equivocal PCS scores of 0 (Fig. 9B). For gene trees with the 20 highest ASTRAL PCS scores for the clade that includes all angiosperms except *Amborella*, all 20 gene trees resolve this clade. ASTRAL PCS scores are therefore consistent with ML concatenation results, the optimal ASTRAL species tree, congruence among independently estimated gene trees, and other support measures (ASTRAL bootstrap, local PP) that show consistent preference for *Amborella* sister to remaining angiosperms.

The distribution of ASTRAL PCS scores lacks prominent outliers (Fig. 9A) in contrast to the distribution of MP-EST PCS scores (Fig. 9B). The highest ASTRAL PCS score for the clade that includes all angiosperms except *Amborella* is +152 quartets (gene tree #52), but this score is just 3% of the total quartet difference between the two conflicting resolutions of *Amborella* (4930 quartets). Indeed, the 39 highest PCS scores for the clade of all angiosperms except *Amborella* total 4884 quartets. Following removal of these 39 high-PCS genes, ASTRAL analysis of the remaining 271 gene trees still supports this clade, but removal of gene trees with the 40 highest PCS scores flips support to the conflicting *Amborella* + *Nuphar* resolution. Unlike MP-EST results, the species tree supported by ASTRAL is highly robust to the removal of many gene trees that strongly favor the preferred topology (Fig. 9B).

Previous reanalyses of the angiosperms dataset suggested that long-branch misplacement of the distant outgroup taxon *Selaginella* might drive the high MP-EST bootstrap support for a species tree in which *Amborella* + *Nuphar* is sister to the remaining angiosperms (Fig. 2D; Simmons and Gatesy, 2015; Simmons, 2017; Simmons et al., accepted). PCS scores help to quantify this bias in misrooted gene trees that position gymnosperms within angiosperms (Fig. 10). Among gene trees with the 20 most negative MP-EST PCS scores for the conflicting *Amborella* + *Nuphar* clade (PCS is negative because MP-EST analysis weakly favors the *Amborella* basal tree; Fig. 9A), 12 gene trees are outliers in which gymnosperms are nested within angiosperms, and just nine of the 20 support the *Amborella* + *Nuphar* clade. In fact, the four gene trees with the most negative MP-EST PCS (gene tree #s 44, 98, 195, 206) contradict the *Amborella* + *Nuphar* clade and instead position *Nuphar* in a basal position relative to *Amborella*, with *Amborella* deeply nested within angiosperms. These four gene trees have little impact on ASTRAL analysis; ASTRAL PCS is 0 for three of these gene trees and slightly negative (−4) for the fourth (Fig. 10). Another five of the 20 gene trees with the most negative MP-EST PCS for *Amborella* + *Nuphar*(gene trees #s 84, 114, 132, 161, 178) also contradict this clade. If the distant outgroup *Selaginella* is pruned from these gene trees, *Amborella* is sister to remaining angiosperms. These five gene trees favor the *Amborella*-basal resolution according to positive ASTRAL PCS scores, which contrasts with the highly negative MP-EST PCS scores for the same gene trees (Fig. 10). PCS scores again demonstrate that different coalescence algorithms can assign very different weights to outlier gene trees (Fig. 10) and that the proportion of negative (conflict) and positive (support) evidence for a clade in the same set of gene trees can vary dramatically between methods (Fig. 9).

**Figure 10.**
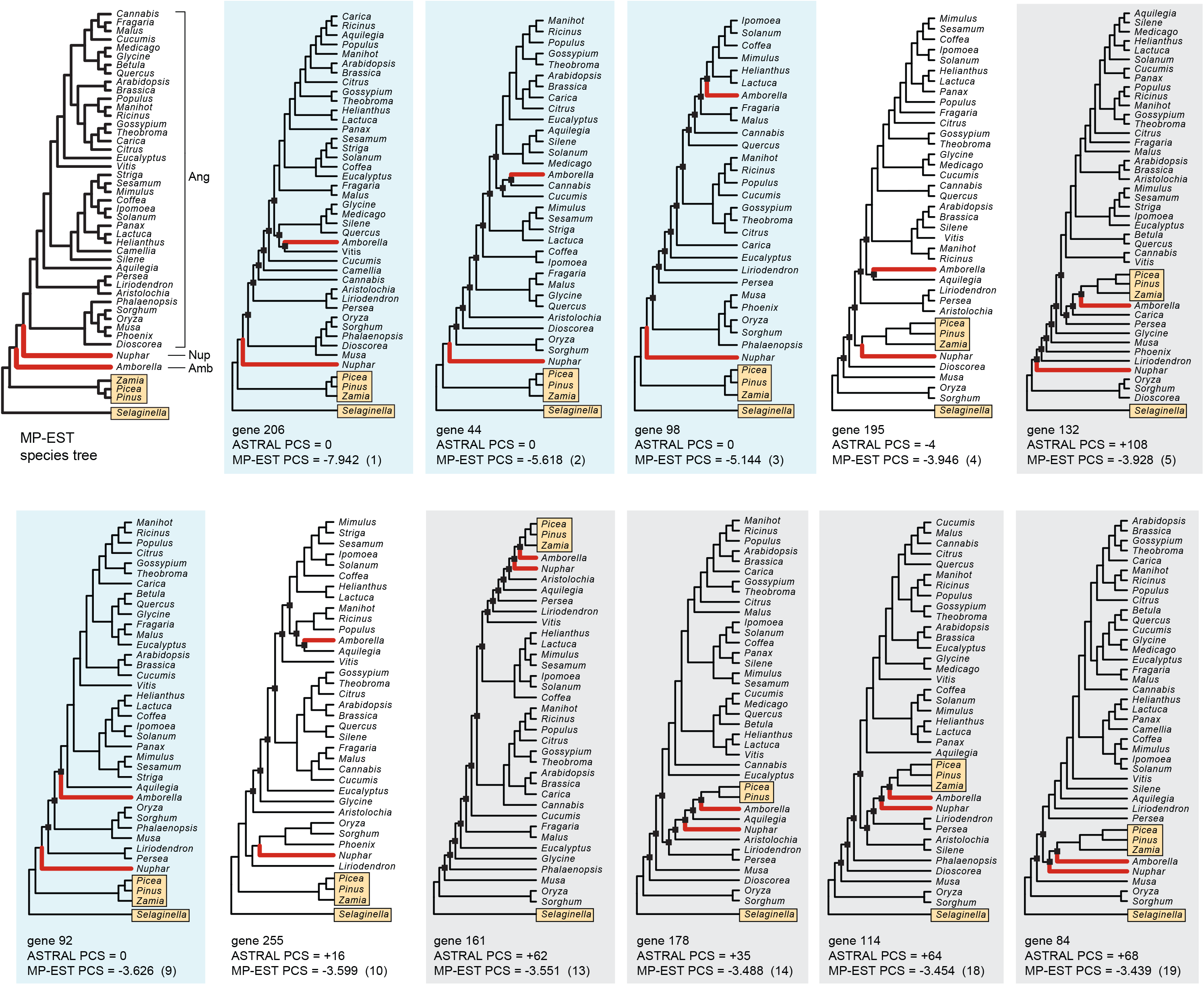
Of the 20 gene trees in the angiosperms dataset (Xi et al., 2014) that most strongly favor the *Amborella* + *Nuphar* clade in MP-EST analysis, eleven contradict the *Amborella* + *Nuphar* clade. These 11 gene trees with very low MP-EST PCS scores also conflict with the optimal MP-EST species tree (top left). The relative rank of each gene tree, in terms of negative MP-EST PCS, is shown in parentheses (e.g., gene tree #132 has the fifth lowest MP-EST PCS score). Six of the 11 gene trees contradict angiosperm monophyly because the distant outgroup *Selaginella* roots on long-branched monocots (*Oryza*, *Sorghum*, *Dioscorea*). Five of these gene trees (light gray backgrounds) have highly negative MP-EST PCS scores but positive ASTRAL PCS scores. In four gene trees where *Nuphar* is basal to *Amborella* (light blue backgrounds), MP-EST PCS is very negative, but ASTRAL PCS is zero. *Amborella* is deeply nested within angiosperms in all 11 gene trees (black squares at nodes). *Amborella* and *Nuphar* are marked by red terminal branches, and outgroup taxa (three gymnosperms and *Selaginella*) are in yellow rectangles. Abbreviations are: Nup = *Nuphar, Amb = Amborella*, Ang = all angiosperms except *Amborella* and *Nuphar.*

If the *Amborella*-basal species tree robustly supported by ASTRAL coalescence and ML concatenation is “right, “ then the high MP-EST coalescence bootstrap support of 99% for the conflicting *Amborella* + *Nuphar* clade must be “wrong.” An explanation for this discrepancy might center on the basic mechanics of the bootstrap procedure when applied in phylogenomic coalescence analyses (Sayyari and Mirarab, 2016; Simmons et al., accepted). Early literature on the bootstrap explicitly warned that this approach should not be applied to small systematic datasets with few characters (Efron, 1979; Felsenstein, 1985, 2004; Sanderson, 1989). The segments of DNA that represent the basic units of coalescence analysis, ‘coalescence genes’ (Hudson, 1990; Doyle, 1992, 1997; Maddison, 1997; Edwards, 2009), can be quite short (Hobolth et al., 2007, 2011) and commonly include few informative characters for short branches (Gatesy and Springer, 2014; Springer and Gatesy, 2016, 2018b). Instead of accounting for gene-tree uncertainty (Liu et al., 2009b, 2015a; Edwards, 2016; Linkem et al., 2016), bootstrapping of nucleotide sites in small coalescence genes actually misrepresents the DNA sequence data collected by the investigator (Sayyari and Mirarab, 2016). In each bootstrap replicate, some nucleotide sites are redundantly sampled while other nucleotide sites in a coalescence gene are completely excluded. Therefore, nodes in gene trees that are supported by single nucleotide substitutions can be lost in the bootstrap replicates (Simmons et al., accepted).

The end result is systematically increased gene-tree heterogeneity (conflicts) in the bootstrap replicates relative to gene trees estimated from the original sequence data (Sayyari and Mirarab, 2016; Simmons et al., accepted). For the angiosperms dataset, Simmons et al. (accepted) documented many more conflicts among gene trees in bootstrap replicates (67.0% of nodes conflict in pairwise comparisons of gene trees) relative to conflicts among gene trees that are based on analyses of the original sequence data (56.0% pairwise incongruence). The same pattern was observed for the amniote dataset (Chiari et al., 2012), which has surprisingly high MP-EST bootstrap support (87-90%) for the controversial crocodilians + turtles clade (Fig. 2A). Conflicts among gene trees in bootstrap replicates (60.9% pairwise incongruence) are higher than conflicts among gene trees that are based on analyses of the original DNA sequence data (51.3% pairwise incongruence). Given that gene-tree-inference error negatively impacts summary coalescence methods (Huang et al., 2010; Knowles et al., 2012; Bayzid and Warnow, 2013; Patel et al., 2013; Lanier et al., 2014), we hypothesize that high MP-EST bootstrap scores for conflicting clades in the amniote dataset (Fig. 2A) and the angiosperms dataset (Fig. 2D) are caused by increased gene-tree-inference errors in bootstrap replicates and biases that are specific to MP-EST (Simmons and Gatesy, 2015; Simmons et al., 2016, accepted; Gatesy et al., 2017).

### 3.2 Common patterns among the four datasets

For the four phylogenomic datasets (Fig. 2), MP-EST and ASTRAL coalescence results disagree at contentious nodes that are central to the main conclusions of each study (Chiari et al., 2012; Zhong et al., 2013; Xi et al., 2014; Linkem et al., 2016). PCS scores provide detailed information on these conflicts between different coalescence methods and a more complete understanding of the support for contested relationships in each study (Figs. 3–10). Several common patterns emerged in these analyses.

#### 3.2.1 Sensitivity to removal of just a few gene trees from genome-scale datasets

In a supermatrix context, measures of phylogenetic stability/sensitivity to the removal of data from analysis were established decades ago and have been executed at the level of taxa (Lanyon, 1985; Siddall, 1995), the most influential characters (Davis, 1993), random subsamples of characters (Penny and Hendy, 1986; Farris et al., 1996; Miller, 2003; Miller and Hormiga, 2004), the most influential genes (Gatesy et al., 1999a, b; Brown and Thomson, 2017; Shen et al., 2017), or random subsamples of genes (Olmstead and Sweere, 1994; Gatesy et al., 1999a, b; Rokas et al., 2003; Narechania et al., 2012). More recently, removals of taxa (Song et al., 2012; Liu et al., 2015b), the most influential gene trees (Gatesy et al., 2017; this study), or random subsamples of gene trees (Song et al., 2012; Simmons and Gatesy, 2015; Edwards, 2016; Richart et al., 2016; Simmons et al., accepted) have been used to assess the stability of species trees inferred using summary coalescence methods.

The logic of stability measures based on data removal is straightforward; the rationale is the same whether taxa, characters, or genes are removed from large datasets. In the most robust datasets, all characters and all genes are congruent with the inferred species tree, many genes have been sampled, and the removal of any character, gene, or taxon from the analysis does not change any relationships in the tree. Furthermore, successively larger deletions of data do not upset the inferred relationships. In the most sensitive datasets, the removal of any character, gene, or taxon overturns all relationships supported by the complete dataset because the number of informative characters is limited, incongruence is rampant, or both. Nearly all empirical datasets are somewhere on the continuum between these two extremes, and stability measures can be used to assess whether a robust sample of data has been collected or whether phylogenetic inferences are more tenuous and hinge on a small percentage of the complete dataset (Gatesy et al., 1999b; Miller, 2003).

Because PCS scores are quantified by the optimality criterion of the coalescence method that is being applied, PCS can be used to rapidly identify gene trees that have a large influence on determining contentious relationships and therefore enable assessments of clade sensitivity to the removal of genes from a phylogenomic dataset (Gatesy et al., 2017). For all four phylogenomic datasets, PCS scores identified high-impact gene trees. Often, the removal of just one to four optimal gene trees from analysis overturned relationships, even for clades with high support according to likelihood ratio tests, the bootstrap, or Bayesian local PP (Figs. 3, 5, 7, 9). These results for species trees parallel recent phylogenomic supermatrix analyses, which showed that controversial clades with high support according to traditional indices (bootstrap or PP) can be overturned by the removal of just a few outlier genes (Brown and Thomson, 2017; Shen et al., 2017). In addition to identifying weaknesses in the support provided by large datasets, PCS scores also can be used to better characterize the robustness of well-supported systematic results based on phylogenomic data. For example, in the ASTRAL analysis of angiosperms, a clade composed of all angiosperms except *Amborella* has local PP of 1.0 (Fig. 2D) and is stable to removal of the 39 gene trees with highest PCS for this clade. This set of genes constitutes 12.6% of the 310 total genes in the dataset (Fig. 9B). By contrast, MP-EST analysis of the same 310 gene trees supports the same clade, but this resolution is not stable to the removal of just one gene tree, 0.3 % of the total (Fig. 9A).

#### 3.2.2 Gene trees with extensive missing taxa are severely downweighted in coalescence analyses

PCS scores quantify the profound impacts of missing taxa on the relative influence of different gene trees in summary coalescence analyses (Fig. 11). Small, partially redundant subtrees (triplets or quartets) summarize the phylogenetic information in gene trees for the MPEST and ASTRAL coalescence methods (Liu et al., 2010; Mirarab et al., 2014). The relationship between gene tree size and potential phylogenetic support is positive, and a single large gene tree that includes thousands of subtrees can overwhelm the support provided by numerous small gene trees that each include just a few subtrees (Figs. 5, 6). Gene trees based on independently segregating regions of non-recombining DNA are the basic units of analysis in coalescence analyses. Therefore, attributing very high weight to some gene trees and very low weight to other gene trees counters the basic logic of phylogenomic coalescence analysis (Gatesy and Springer, 2014). Simulation studies have addressed the effects of missing data in species-tree analysis and have noted decreased phylogenetic accuracy in some hypothetical contexts (Vachaspati and Warnow, 2015; Xi and Davis, 2016; Sayyari et al., 2017; Molloy and Warnow, 2018). Likewise, it is evident from equations in the original descriptions of MP-EST (Liu et al., 2010) and ASTRAL (Mirarab et al., 2014) that, all else being equal, large gene trees are assigned more weight than small gene trees.

**Figure 11.**
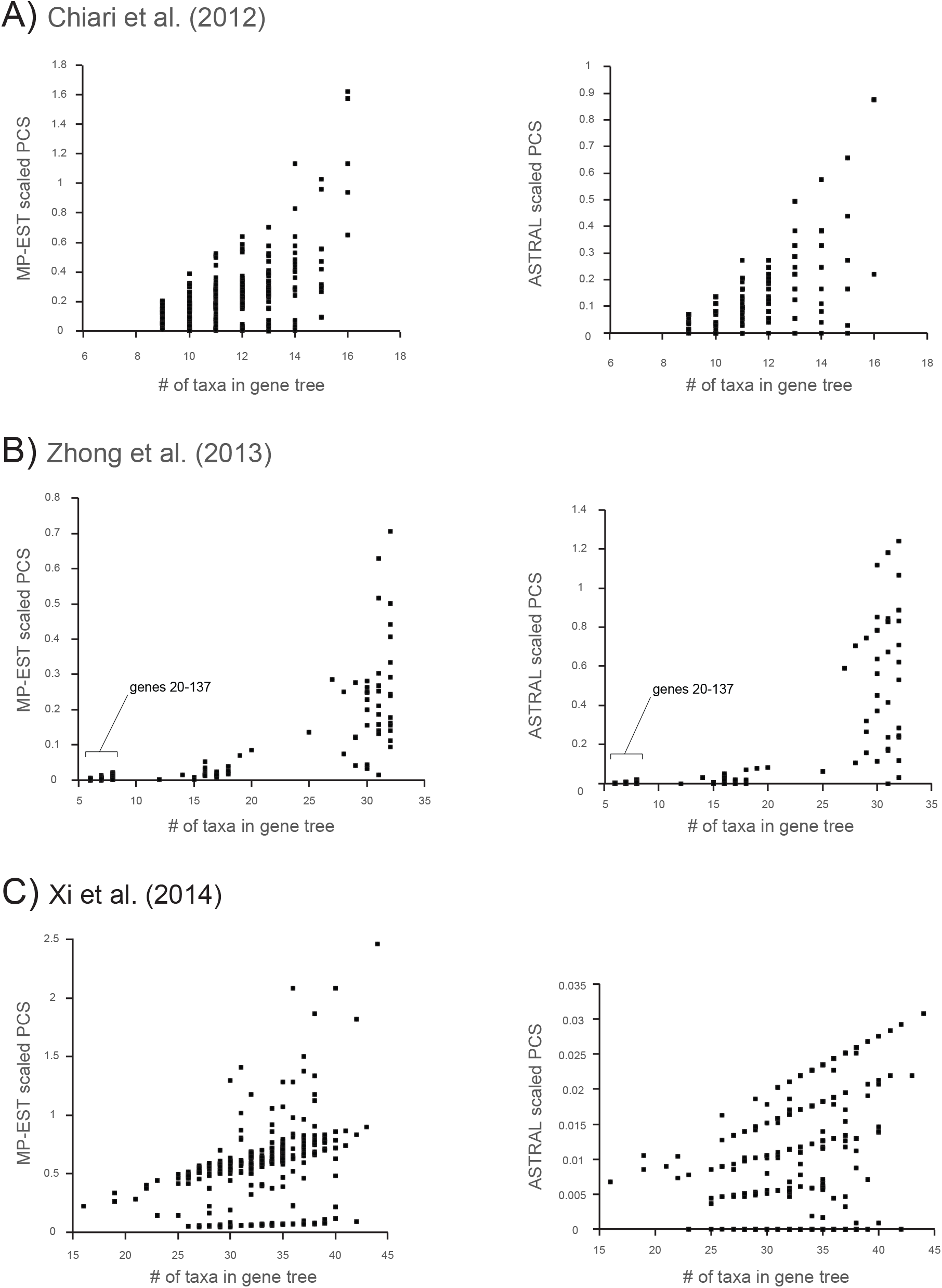
Gene tree size (number of taxa) versus the absolute value of scaled PCS for three phylogenomic datasets with missing taxa in some gene trees: A) Chiari et al. (2012), B) Zhong et al. (2013), and C) Xi et al. (2014). For both MP-EST (left) and ASTRAL (right), the absolute values of scaled PCS scores (Y axis) for contentious clades (Fig. 2A-C) are plotted against gene tree size (X axis). Note that the scaled PCS scores of small gene trees are consistently lower than scaled PCS scores of large gene trees for all three datasets; gene tree size defines the maximum influence that a gene tree can have at a particular node. The set of 118 very small gene trees (each 6-8 taxa; gene trees #20-137) from Zhong et al. (2013) that have little influence in MP-EST and ASTRAL analyses are bracketed.

For any particular empirical dataset, however, PCS scores directly quantify how missing taxa in gene trees directly impact support for any given clade. PCS for the land-plants dataset (Zhong et al., 2013) provides a compelling example because of missing taxa in numerous gene trees (Figs. 5–6). Similar, but less dramatic effects of gene-tree size are apparent in the amniote and angiosperm datasets. For all datasets with missing taxa in this study, gene trees with more taxa have dramatically greater influence in MP-EST and ASTRAL coalescence analyses than do gene trees with fewer taxa (Fig. 11). Many taxa in a gene tree do not guarantee extremely high or extremely low PCS for a particular clade. PCS can be zero for a small gene tree or for a large gene tree depending on relationships in the gene tree, however, larger gene trees have more *potential* for high PCS. Small gene trees have a limited number of unique triplets or quartets, and PCS scores are tightly bounded by this constraint. By contrast, large gene trees can have a much broader range of PCS scores (Figs. 4, 6, 11), simply because there are many more relevant triplets and quartets in gene trees with many taxa.

Genes with better taxonomic sampling might, on average, yield more accurate gene trees (Zwickl and Hillis, 2002; Hedtke et al., 2006), but large gene trees should not be upweighted in proportion to the number of quartets or triplets in a gene tree that are informative for a given clade (Fig. 11). The supermatrix approach (Kluge, 1989; Nixon and Carpenter, 1996) has been criticized for misweighting data because independence of each nucleotide site (= character) is assumed, and the close linkage of some sites (and associated dependence) is not accounted for in a biologically realistic way (Kubatko and Degnan, 2007; Edwards, 2009; Liu and Edwards, 2009; Edwards et al., 2016). For example, a large, effectively non-recombining locus that is rapidly evolving, such as mitochondrial DNA in animals, might overwhelm support from many independently sorting nuclear loci in a combined supermatrix (McVay and Carstens, 2013). Gross under-weighting of gene trees with extensive missing taxa could have even more devastating effects in summary coalescence analyses. For the land-plants dataset, 118 small gene trees (#s 20-137) are practically ignored in ASTRAL coalescence analyses, and final systematic results are driven by the 66 larger gene trees (Figs. 5B, 11). Most phylogenomic datasets contain less missing data, but even subtle differences in taxon sampling can impact results (Fig. 11). Support for one clade over a conflicting clade might tip one way or another simply because a particular resolution is supported by a few more large gene trees, rather than the majority of independently segregating loci (Figs. 3A, 5). Because gene trees are decomposed into overlapping sets of subtrees in MP-EST and ASTRAL, both methods are subject to this effect when missing taxa are present (Fig. 11). PCS scores help to characterize the instability of a clade that is driven by different gene-tree sizes when these species-tree methods are applied.

PCS does not correct distortions in phylogenetic results due to missing taxa. A correction for missing-data effects would require an adjustment in how quartets or triplets from gene trees of unequal sizes are weighted in summary coalescence analyses using MP-EST or ASTRAL. Alternatively, coalescence methods that do not break gene trees down into constituent triplets or quartets could be applied (e.g., ‘minimize deep coalescence’ – Maddison, 1995; *BEAST – Heled and Drummond, 2010), or analysis could be restricted to gene trees with no missing taxa (e.g., Song et al., 2012; Linkem et al., 2016).

#### 3.2.3 Overweighting of outlier gene trees in MP-EST coalescence analyses

MP-EST and ASTRAL coalescence methods do not treat gene trees as supporting evidence in the same way (Fig. 12). For MP-EST, gene trees are decomposed into rooted triplets, and inferred branch lengths in species trees influence the pseudo-likelihood score that is used to choose among different species tree hypotheses (Liu et al., 2010). For ASTRAL, gene trees are instead summarized by unrooted quartets, and branch lengths have no impact on the optimality score, which is the number of quartets in gene trees that are compatible with different species-tree topologies (Mirarab et al., 2014). Previous work has revealed situations (e.g., Figs. 1, 2) where MP-EST and ASTRAL analyses of the same set of gene trees yield different species trees when there are many misrooted gene trees (Simmons and Gatesy, 2015), numerous conflicts among gene trees (Mirarab and Warnow, 2015; Springer and Gatesy, 2016), or displacements of ingroup taxa to very basal positions in large gene trees (Gatesy et al., 2017). These conflicts between methods may be driven by branch-length estimations in MP-EST species trees (Mirarab and Warnow, 2015), differences in how rooted triplets and unrooted quartets summarize gene trees (Simmons and Gatesy, 2015; Gatesy et al., 2017), homology problems (Springer and Gatesy, 2016, 2018a, c; Gatesy and Springer, 2017), or inadequate tree searches (Springer and Gatesy, 2016; Gatesy et al., 2017).

**Figure 12.**
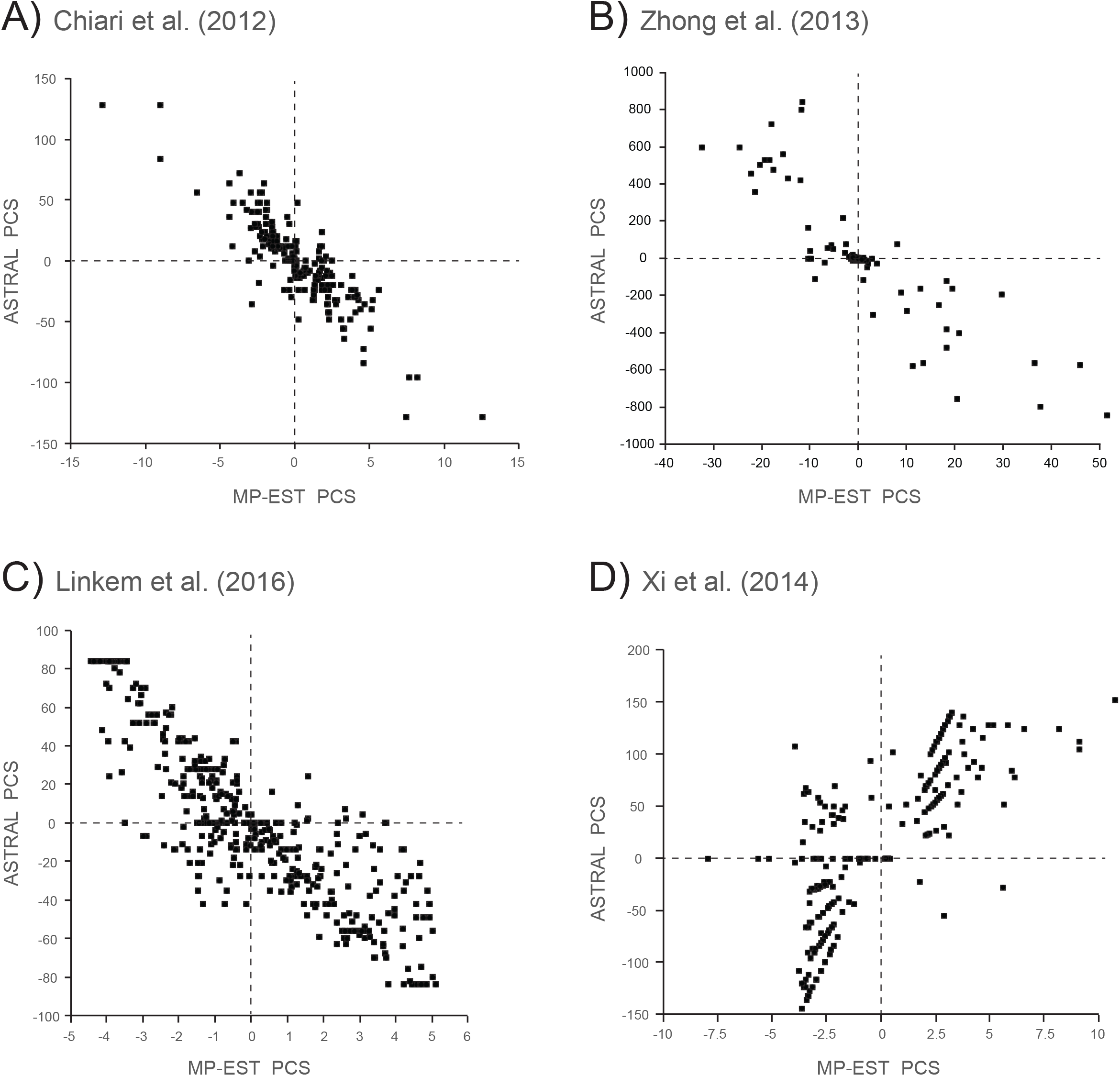
A comparison of MP-EST PCS scores versus ASTRAL PCS scores for four phylogenomic datasets: A) Chiari et al. (2012), B) Zhong et al. (2013), C) Linkem et al. (2016), and D) Xi et al. (2014). For each gene tree, MP-EST PCS (X axis) is plotted against ASTRAL PCS (Y axis) at contentious nodes (Fig. 2). For three datasets (A-C), MP-EST and ASTRAL support alternative phylogenomic resolutions (Fig. 2A-C). For the angiosperms dataset (Xi et al., 2014), bootstrap support is high for conflicting MP-EST and ASTRAL resolutions (Fig. 2D), but both methods support *Amborella* as sister to remaining angiosperms in analyses of optimal gene trees (Fig. 9). In the plot of MP-EST PCS versus ASTRAL PCS for Xi et al. (2014), note that the three gene trees with the most negative MP-EST PCS each have ASTRAL PCS scores of zero (see Fig. 10). Scatter in the four plots indicates discrepancies in how gene trees are translated into support for alternative species trees when different summary coalescence methods are applied (Fig. 2).

Because PCS is quantified by the specific optimality criterion of the coalescence method that is being applied, PCS scores for a clade effectively discern the most influential gene trees for that coalescence method, i.e., gene trees that strongly favor the supported clade versus an alternative species tree that lacks the clade. Therefore, when MP-EST and ASTRAL are applied to the same set of optimal gene trees, PCS scores clearly display differences in how the alternative methods interpret gene trees as evidence for competing phylogenetic hypotheses (Fig. 12). For the datasets examined here, outlier gene trees commonly provide the highest MP-EST scores for controversial clades that conflict with ASTRAL- and supermatrix-based inferences (Figs. 3, 7, 9), and these outlier gene trees with very high MP-EST PCS oftentimes contradict the contentious clade (Figs. 8, 10). By contrast, the highest ASTRAL PCS scores for contested clades come from gene trees that actually resolve the clade of interest. For the angiosperms dataset, the misrooting artifact noted by Simmons and Gatesy (2015) is evident in five gene trees with highly negative MP-EST PCS scores (Fig. 10). All of these gene trees (#s 84, 114, 132, 161, 178) actually contradict the controversial *Amborella* + *Nuphar* clade, but many rooted triplets in these gene trees *do* support *Amborella* + *Nuphar*. The same cannot be said for unrooted quartets, which are the currency of ASTRAL analysis. More unrooted quartets are instead compatible with the *Amborella*-basal tree than with the *Amborella* + *Nuphar* tree, and if the distant *Selaginella* outgroup is pruned away, these five gene trees *do* support *Amborella* as basal when root position is determined by the less divergent gymnosperms clade (Fig. 10).

Another MP-EST bias is evident in PCS scores for the angiosperms, land plants, and skinks datasets. For each dataset, many of the gene trees with high MP-EST PCS scores for the conflicting clade resolve the basal lineage of that clade as nested well within its putative sister group (Figs. 6, 8, 10). This ‘apical nesting bias’ has no impact on ASTRAL PCS scores (e.g., gene tree #174 vs. gene tree #243; Fig. 8). In many cases, the aberrant gene trees with high MPEST PCS do not even support the clade of interest, and the ‘apical nesting bias’ can occur with (Fig. 10) or without (Fig. 8) missing taxa in a phylogenomic dataset. The resolution of a lineage well within its sister group does not increase counts of compatible triplets or quartets; there is no difference in the fit of these small subtrees in gene trees to the alternative species-tree topologies. We could not rationalize any coherent justification, in the context of the multispecies coalescent, for giving such high relative weights to these aberrant gene-tree topologies. PCS scores have helped discern a ‘basal dragdown bias’ in MP-EST (Figs. 1, 9; Gatesy et al., 2017) as well as an ‘apical nesting bias’ in MP-EST (Figs. 5, 8, 10). These two biases, which are in opposite directions for a given species tree, hint at a single, more general, underlying bias that merits further investigation.

#### 3.2.4 Logically consistent support indices for phylogenomic coalescence analyses

Recent work has highlighted problems with resampling nucleotide sites when bootstrapping genes in summary coalescence analyses (Sayyari and Mirarab, 2016; Simmons et al., accepted). This issue has provided the impetus for the development of support measures that do not require resampling sites with replacement. PCS is based on analysis of optimal gene trees, does not require resampling of nucleotide sites, is logically consistent with the optimality criteria of commonly-used coalescence methods, and can be used to assess the stability of a clade to the removal of outlier gene trees (Gatesy et al., 2017). More recently, Simmons et al. (accepted) argued for the utility of resampling gene trees instead of sites in bootstrap or jackknife analyses in a summary coalescent context.

Sayyari and Mirarab (2016) used counts of quartets in optimal gene trees for conflicting species-tree resolutions to derive a Bayesian measure of support—Bayesian local PP. Each optimal gene tree for a dataset is given equal weight in this approach, and a local PP can be generated rapidly for each clade in a species tree as well two alternative suboptimal resolutions for each clade (Zhang et al., 2017). Like the initial formulation of PCS (Gatesy et al., 2017), the local PP focuses on comparisons between a supported clade and single local branch swaps relative to the supported clade. Because the Bayesian local PP is implemented in ASTRAL (Mirarab and Warnow, 2015) and is computed rapidly, this statistic commonly is reported for ASTRAL species trees. Unlike CS and PCS, which are compatible with the optimality criterion of ASTRAL, however, the local PP can show high Bayesian support for a clade that is contradicted by the optimal ASTRAL species tree. This is because quartets are the basic units of analysis in ASTRAL (Mirarab et al., 2014), while individual gene trees are the basic units of analysis for the local PP (Sayyari and Mirarab, 2016). In datasets where many taxa are missing and/or conflicts among gene trees are extensive (e.g., Chiari et al., 2012; Zhong et al., 2013), this inconsistency can be a factor when interpreting local PPs (Figs. 3, 5).

Taxonomic congruence approaches (Nelson, 1979; Miyamoto and Fitch, 1995) that assess support for a clade via agreements and conflicts with independently estimated gene trees (Salichos and Rokas, 2013; Salichos et al., 2014; Smith et al., 2015; Kobert et al., 2016) also are inconsistent with the underpinnings of summary coalescence methods such as MP-EST and ASTRAL. This is because even highly incongruent gene trees that conflict with all three resolutions at a node can contribute relatively more support to one of the three alternative resolutions (e.g., Fig. 10). Indeed, a clade supported by a summary coalescence analysis might not be found in *any* of the input gene trees, thus indicating that the clade is supported only by emergent ‘hidden support’ (Gatesy and Springer, 2014; Gatesy et al., 2017). A taxonomic congruence approach would indicate no support for that clade from *any independently-estimated* gene tree, even though the clade is favored by the overall coalescence analysis of optimal gene trees. The conclusion that there is no support for a supported clade is due to the fact that the optimality criterion of the coalescence method applied (e.g., the fit of unrooted quartets to different species trees determines the optimal species tree) is at odds with the logical basis of the support index (the number of times that a clade is corroborated by independently-estimated gene trees determines the level of support). An argument can therefore be made for measures of phylogenetic support that are compatible with the basic mechanics of the phylogenetic methods that are applied. Examples include Bremer support for parsimony analysis (Bremer, 1994), likelihood ratio tests for ML supermatrix analysis (Huelsenbeck et al., 1996), jackknifing of optimal gene trees for summary coalescence methods (Simmons et al., accepted), and PPs for Bayesian supermatrix (Nylander et al., 2004) or coalescence analyses (Heled and Drummond, 2010).

## 4. Conclusions

1. PCS quantifies the positive or negative influence of each gene tree in a phylogenomic dataset for clades supported by summary coalescence methods such as MP-EST and ASTRAL that employ optimality criteria to estimate species trees.
2. PCS can be generalized to comparisons between any two conflicting species trees, in the same manner as partitioned support for clades supported by supermatrices (Gatesy et al., 1999a; Shen et al., 2017).
3. When two conflicting species trees are compared, ‘scaled PCS’ summarizes the proportional contribution (positive, negative, or zero) of each gene tree to the total coalescence support (CS) for a clade.
4. Automation of PCS permits rapid calculation of PCS scores for genome-scale datasets of hundreds or thousands of gene trees using either MP-EST or ASTRAL (see pcs_mpest and pcs_astral repositories at https://github.com/dbsloan).
5. Application of PCS to empirical phylogenomic datasets (Chiari et al., 2012; Zhong et al., 2013; Xi et al., 2014; Linkem et al., 2016) shows that, in all four cases, contentious relationships that were central to the primary conclusions of the original studies are sensitive to the removal of just one to four gene trees with high PCS scores (Figs. 3, 5, 7, 9).
6. In some cases, clades with high support scores according to likelihood ratio tests (*P < 0.05)*, Bayesian local PP (1.0), or the bootstrap (90%, 99%) flip to alternative phylogenomic resolutions with the removal of just one or two overweighted gene trees that were identified by PCS (Figs. 3, 5A, 9A).
7. PCS scores quantify the striking impact of missing taxa in gene trees for both MP-EST and ASTRAL coalescence analyses at contested nodes (Figs. 3–6, 11). For the land-plants dataset (Zhong et al., 2013), one large gene tree is more influential than >100 small gene trees in ASTRAL analysis due to discrepant counts of informative quartets in big versus small gene trees (Figs. 5, 6). Attributing exceptionally high weight to some gene trees and very low weight to other gene trees counters the basic logic of phylogenomic coalescence analysis (Gatesy and Springer, 2014).
8. PCS scores quantify the overweighting of outlier gene trees in MP-EST analysis at critical nodes. Some of these outliers imply deep coalescences (retentions of ancestral polymorphism) that are not credible (Figs. 3, 4, 9A, 10). Furthermore, gene trees that do not even support the clade of interest can have extremely high MP-EST PCS scores for that clade (Figs. 8, 10). ‘Misrooting bias’ (Simmons and Gatesy, 2015; Fig. 10), ‘basal-dragdown bias’ (Gatesy et al., 2017; Figs. 1, 9A), and ‘apical-nesting bias’ (Figs. 5, 8, 10), which are revealed by MP-EST PCS, might explain incongruence between MP-EST and ASTRAL coalescence results in some cases.
9. PCS scores are consistent with the optimality criterion of the coalescence method that is applied, and because optimal gene trees are used to compute PCS, extraneous conflict among gene trees that is caused by bootstrap resampling of sites (Sayyari and Mirarab, 2016; Simmons et al., accepted) is not a factor. Bayesian local PPs are based on optimal gene trees (Sayyari and Mirarab, 2016), but because the assumptions of this support measure are not completely compatible with the optimality criterion of ASTRAL, PPs of 0.00 for supported clades and PPs of 1.00 for contradicted clades are possible (Fig. 3).
10. The development of partitioned support measures for distance-based coalescence methods (STAR, STEAC, NJ-ST, ASTRID: Liu et al., 2009b, Liu and Yu, 2011; Vachaspati and Warnow, 2015) should be a priority to make sense of conflicting relationships favored by different phylogenomic coalescence methods (e.g., Esselstyn et al., 2017; Gatesy and Springer, 2017; Liu et al., 2017).

## 5. Acknowledgements

We thank A. de Queiroz, C. Hayashi, and B. Smith for providing helpful comments on various drafts of the manuscript. Discussions with C. Ané, S. Mirarab, and T. Warnow provided clarity on various aspects of summary coalescence analysis. F. Delsuc provided useful information on the amniotes dataset. This research was funded by NSF grant DEB-1457735 (J.G. and M.S.S.), NSF grants MCB-1733227 and IOS-1829176 (D.B.S), and an NSF-funded GAUSSI Graduate Fellowship (DGE-1450032) to J.M.W.

## Abbreviations

ASTRAL: accurate species tree algorithm
ASTRID: accurate species trees from internode distances
ILS: incomplete lineage sorting
ML: maximum likelihood
MP-EST: maximum pseudo-likelihood for estimating species trees
MY: million years
NJ-ST: neighbor joining species tree
PCS: partitioned coalescence support
PP: posterior probability
STAR: species tree estimation using average ranks of coalescences
STEAC: species tree estimation using average coalescence times
UCE: ultraconserved element

## References

Arcila, D., Ortí, G., Vari, R., Armbruster, J.W., Stiassny, M.L.J., Ko, K.D., Sabaj, M.H., Lundberg, J., Revell, L.J., Betancur-R, R., 2017. Genome-wide interrogation advances resolution of recalcitrant groups in the tree of life. Nature Ecol. Evol. 1, 0020.

Baker, R.H., DeSalle, R., 1997. Multiple sources of character information and the phylogeny of Hawaiian drosophilids. Syst. Biol. 46, 654–673.

Barrett M., Donoghue M.J., Sober E., 1991. Against consensus. Syst. Zool. 40, 486–493.

Bayzid, M.S., Warnow, T., 2013. Naive binning improves phylogenomic analyses. Bioinformatics 29, 2277–2284.

Bayzid, M.S., Mirarab, S., Boussau, B., Warnow, T., 2015. Weighted statistical binning: enabling statistically consistent genome-scale phylogenetic analyses. PLoS ONE 10, e0129183.

Bell, C.D., Soltis, D.E., Soltis, P.S., 2010. The age and diversification of the angiosperms re-revisited. Amer. J. Bot. 97, 1296–1303.

Bremer, K., 1994. Branch support and tree stability. Cladistics 10, 295–304.

Brown, J.M., Thomson, R.C., 2017. Bayes factors unmask highly variable information content, bias, and extreme influence in phylogenomic analyses. Syst. Biol. 66, 517–530.

Chiari, Y., Cahais, V., Galtier N., Delsuc, F., 2012. Phylogenomic analyses support the position of turtles as the sister group of birds and crocodiles (Archosauria). BMC Biol. 10, 65.

Chippindale, P., Wiens, J., 1994. Weighting, partitioning, and combining characters in phylogenetic analysis. Syst. Biol. 43, 278–287.

Davis, J., 1993. Character removal as a means for assessing stability of clades. Cladistics 9, 201–210.

Degnan, J.H., Rosenberg, N.A., 2006. Discordance of species trees with their most likely gene trees. PLoS Genetics 2, e68.

de Queiroz, A., Gatesy, J., 2007. The supermatrix approach to systematics. Trends Ecol. Evol. 22, 34–41.

Doyle, J.J., 1992. Gene trees and species trees: molecular systematics as one-character taxonomy. Syst. Bot. 17, 144–163.

Doyle, J.J., 1997. Trees within trees: genes and species, molecules and morphology. Syst. Biol. 46, 537–553.

Edwards, S.V. 2009. Is a new and general theory of molecular systematics emerging? Evolution 63, 1–19.

Edwards, S.V., 2016. Phylogenomic subsampling: a brief review. Zool. Scr. 45, 63–74.

Edwards, S.V., Xi, Z., Janke, A., Faircloth, B.C., McCormack, J.E., Glenn, T.C., Zhong, B., Wu, S., Lemmon, E.M., Lemmon, A.R., Leaché, A.D., Liu, L., Davis, C.C., 2016. Implementing and testing the multispecies coalescent model: a valuable paradigm for phylogenomics. Mol. Phylogenet. Evol. 94, 447–462.

Efron, B., 1979. Bootstrap methods: another look at the jackknife. Ann. Stat. 7, 1–26.

Esselstyn, J.A., Oliveros, C.H., Swanson, M.T., Faircloth, B.C., 2017. Investigating difficult nodes in the placental mammal tree with expanded taxon sampling and thousands of ultraconserved elements. Genome Biol. Evol. 9, 2308–2321.

Farris, J.S., Albert, V.A., Källersjö, M., Lipscomb, D., Kluge, A.G., 1996. Parsimony jackknifing outperforms neighbor-joining. Cladistics 12, 99–124.

Felsenstein, J., 1985. Confidence limits on phylogenies: an approach using the bootstrap. Evolution 39, 783–791.

Felsenstein, J., 2004. Inferring phylogenies. Sunderland, Mass., Sinauer Associates, Inc.

Gatesy J., Baker, R.H., 2005. Hidden likelihood support in genomic data: can forty-five wrongs make a right? Syst. Biol. 54, 483–492.

Gatesy, J., Springer, M.S., 2014. Phylogenetic analysis at deep timescales: unreliable gene trees, bypassed hidden support, and the coalescence/concatalescence conundrum. Mol. Phylogenet. Evol. 80, 231–266.

Gatesy, J., Springer, M.S., 2017. Phylogenomic red flags: homology errors and zombie lineages in the evolutionary diversification of placental mammals. Proc. Natl. Acad. Sci. USA 114, E9431–E9432.

Gatesy, J., O’Grady, P., Baker, R.H., 1999a. Corroboration among data sets in simultaneous analysis: hidden support for phylogenetic relationships among higher level artiodactyl taxa. Cladistics 15, 271–313.

Gatesy, J., Milinkovitch, M.C., Waddell, V., Stanhope, M., 1999b. Stability of cladistic relationships between Cetacea and higher-level artiodactyl taxa. Syst. Biol. 48, 6–20.

Gatesy, J., Meredith, R.W., Janečka, J.E., Simmons, M.P., Murphy, W.J., Springer, M.S., 2017. Resolution of a concatenation/coalescence kerfuffle: partitioned coalescence support and a robust family-level tree for Mammalia. Cladistics 33, 295–332.

Hedtke, S.M., Townsend, T.M., Hillis, D.M., 2006. Resolution of phylogenetic conflict in large data sets by increased taxon sampling. Syst. Biol. 55, 522–529.

Heled, J., Drummond, A.J., 2010. Bayesian inference of species trees from multilocus data. Mol. Biol. Evol. 27, 570–580.

Hobolth, A., Christensen, O.F., Mailund, T., Schierup, M.H., 2007. Genomic relationships and speciation times of human, chimpanzee, and gorilla inferred from a coalescent hidden Markov model. PLoS Genet. 3, e7.

Hobolth, A., Dutheil, J.Y., Hawks, J., Schierup, M.H., Mailund, T., 2011. Incomplete lineage sorting patterns among human, chimpanzee, and orangutan suggest recent orangutan speciation and widespread selection. Genome Res. 21, 349–356.

Hosner, P.A., Faircloth, B.C., Glenn, T.C., Braun, E.L., Kimball. R.T., 2016. Avoiding missing data biases in phylogenomic inference: an empirical study in the landfowl (Aves: Galliformes). Mol. Biol. Evol. 33, 1110–1125.

Hovmöller, R., Knowles, L.L., Kubatko, L.S., 2013. Effects of missing data on species tree estimation under the coalescent. Mol. Phylogenet. Evol. 69: 1057–1062.

Huang, H., He, Q., Kubatko, L.S., Knowles, L.L., 2010. Sources of error inherent in species-tree estimation: impact of mutational and coalescent effects on accuracy and implications for choosing among different methods. Syst. Biol. 59, 573–583.

Hudson, R.R., 1990. Gene genealogies and the coalescent process. Oxford Surv. Evol. Biol. 7, 1–44.

Huelsenbeck, J.P., Hillis, D.M., Nielsen, R., 1996. A likelihood ratio test of monophyly. Syst. Biol. 45, 546–548.

Jarvis, E.D., Mirarab, S., Aberer, A.J., Li, B., Houde, P., Li, C., Ho, S.Y.W., Faircloth, B.C., Nabholz, B., Howard, J.T., et al., 2014. Whole-genome analyses resolve early branches in the tree of life of modern birds. Science 346, 1320–1331.

Jeffroy, O., Brinkmann, H., Delsuc, F., Philippe, H., 2006. Phylogenomics: the beginning of incongruence? Trends. Genet. 22, 225–231

Kluge, A.G., 1989. A concern for evidence and a phylogenetic hypothesis for relationships among *Epicrates* (Boidae, Serpentes). Syst. Zool. 38, 7–25.

Knowles, L.L., Lanier, H.C., Kilmov, P., He, Q., 2012. Full modeling versus summarizing phylogenetic uncertainty: method choice and species-tree accuracy. Mol. Phylogenet. Evol. 65, 501–509.

Kobert, K., Salichos, L., Rokas, A., Stamatakis, A., 2016. Computing the internode certainty and related measures from partial gene trees. Mol. Biol. Evol. 33, 1606–1617.

Kubatko, L.S., Degnan, J.H., 2007. Inconsistency of phylogenetic estimates from concatenated data under coalescence. Syst. Biol. 56, 17–24.

Lanier, H.C., Huang, H., Knowles, L.L., 2014. How low can you go? The effects of mutation rate on the accuracy of species-tree estimation. Mol. Phylogenet. Evol. 70, 112–119.

Lanyon, S., 1985. Detecting internal inconsistencies in distance data. Syst. Zool. 34, 397–403.

Lee, M., Hugall, A., 2003. Partitioned likelihood support and the evaluation of data set conflict. Syst. Biol. 52, 15–22.

Leffler, E., Gao, Z., Pfeifer, S., Ségurel, L., Auton, A., Venn, O., Bowden, R., Bontrop, R., Wall, J., Sella, G., Donnelly, P., McVean, G., Przeworski, M., 2013. Multiple instances of ancient balancing selection shared between humans and chimpanzees. Science 339, 1578–1582.

Linkem, C.W., Minin, V.N., Leaché, A.D., 2016. Detecting the anomaly zone in species trees and evidence for a misleading signal in higher-level skink phylogeny (Squamata: Scincidae). Syst. Biol. 65, 465–477.

Liu, L., Edwards, S.V., 2009. Phylogenetic analysis in the anomaly zone. Syst. Biol. 58, 452–460.

Liu, L., Yu, L., Kubatko, L.S., Pearl, D.K., Edwards, S.V., 2009a. Coalescent methods for estimating phylogenetic trees. Mol. Phylogenet. Evol. 53, 320–328.

Liu, L., Yu, L., Pearl, D.K., Edwards, S.V., 2009b. Estimating species phylogenies using coalescence times among sequences. Syst. Biol. 58, 468–477.

Liu, L., Yu, L., Edwards, S.V., 2010. A maximum pseudo-likelihood approach for estimating species trees under the coalescent model. BMC Evol. Biol. 10, 302.

Liu, L., Yu, L., 2011. Estimating species trees from unrooted gene trees. Syst. Biol. 60, 661–667.

Liu, L., Xi, Z., Wu, S., Davis, C.C., Edwards, S.V., 2015a. Estimating phylogenetic trees from genome-scale data. Ann. NY Acad. Sci. 1360, 36–53.

Liu, L., Xi, Z., Davis, C.C., 2015b. Coalescent methods are robust to the simultaneous effects of long branches and incomplete lineage sorting. Mol. Biol. Evol. 32, 791–805.

Liu, L., Zhang, J., Rheindt, F.E., Lei, F., Qu, Y., Wang, Y., Sullivan, C., Ni, W., Wang. J., Yang. F., Chen, J., Edwards, S.V., Meng, J., Wu, S., 2017. Genomic evidence reveals a radiation of placental mammals uninterrupted by the KPg boundary. Proc. Natl. Acad. Sci. USA 114, E7282–E7290.

Maddison, W.P., 1997. Gene trees in species trees. Syst. Biol. 46, 523–536.

Magallón, S., Hilu, K.W., Quandt, D., 2013. Land plant evolutionary timeline: gene effects are secondary to fossil constraints in relaxed clock estimation of age and substitution rates. Amer. J. Bot. 100, 556–573.

McVay, J., Carstens, B., 2013. Phylogenetic model choice: justifying a species tree or concatenation analysis. J. Phylogenet. Evol. Biol. 1, 114.

Mendes, F.K.. Hahn, M.W., 2018. Why concatenation fails near the anomaly zone. Syst. Biol. 67, 158–169.

Meredith, R.W., Janecka, J.E., Gatesy, J., Ryder, O.A., Fisher, C.A., Teeling, E.C., Goodbla, A., Eizirik, E., Simão, T.L.L., Stadler, T., Rabosky, D.L., Honeycutt, R.L., Flynn, J.J., Ingram, C.M., Steiner, C., Williams, T.L., Robinson, T.J., Burk-Herrick, A., Westerman, M., Ayoub, N.A., Springer, M.S., Murphy, W.J., 2011. Impacts of the Cretaceous terrestrial revolution and KPg extinction on mammal diversification. Science 334, 521–524.

Miller, J.A., 2003. Assessing progress in systematics with continuous jackknife function analysis. Syst. Biol. 52: 55–65.

Miller, J.A., Hormiga, G., 2004. Clade stability and the addition of data: a case study from erigonine spiders (Araneae: Linyphiidae, Erigoninae). Cladistics 20, 385–442.

Mirarab, S., Warnow, T., 2015. ASTRAL-II: coalescent-based species tree estimation with many hundreds of taxa and thousands of genes. Bioinformatics 31, 144–152.

Mirarab, S., Reaz, R., Bayzid, M. S., Zimmermann, T., Swenson, M.S., Warnow, T., 2014. ASTRAL: genome-scale coalescent-based species tree estimation. Bioinformatics 30, 1541–1548.

Mirarab, S., Bayzid, M.S., Warnow, T., 2016. Evaluating summary methods for multilocus species tree estimation in the presence of incomplete lineage sorting. Syst. Biol. 65, 366–380.

Miyamoto, M., 1985. Consensus cladograms and general classifications. Cladistics 1, 186–189.

Miyamoto, M.M., Fitch, W.M., 1995. Testing species phylogenies and phylogenetic methods with congruence. Syst. Biol. 44, 64–76.

Molloy, E.K., Warnow, T., 2018. To include or not to include; the impact of gene filtering on species tree estimation methods. Syst. Biol. 67, 285–303.

Narechania A, Baker, R.H., Sit, R., Kolokotronis, S., DeSalle, R., Planet, P.J., 2012. Random addition concatenation analysis: a novel approach to the exploration of phylogenomic signal reveals strong agreement between core and shell genomic partitions in the cyanobacteria. Genome Biol. Evol. 4, 30–43.

Nelson, G., 1979. Cladistic analysis and synthesis: principles and definitions, with a historical note on Adanson’s *Familles des plantes* (1763-1764). Syst. Zool. 28, 1–21.

Nixon, K.C., Carpenter, J.M., 1996. On simultaneous analysis. Cladistics 12, 221–242.

Nylander, J.A.A., Ronquist, F., Huelsenbeck, J.P., Nieves-Aldrey, J.L., 2004. Bayesian phylogenetic analysis of combined data. Syst. Biol. 53, 47–67.

Olmstead, R., Sweere, J., 1994. Combining data in phylogenetic systematics: an empirical approach using three molecular data sets in the Solanaceae. Syst. Biol. 43, 467–481.

Patel, S., Kimball, R.T., Braun, E.L., 2013. Error in phylogenetic estimation for bushes in the tree of life. Phylogenet. Evol. Biol. 1, 110.

Penny, D., Hendy, M., 1986. Estimating the reliability of evolutionary trees. Mol. Biol. Evol. 3, 403–417.

Philippe, H., de Vienne, D.M., Ranwez, V., Roure, B., Baurain, D., Delsuc, F., 2017. Pitfalls in supermatrix phylogenomics. Eur. J. Taxon. 283, 1–25.

Piertney, S.B., Oliver, M.K., 2006. The evolutionary ecology of the major histocompatibility complex. Heredity 96, 7–21

Richart, C.H., Hayashi, C.Y., Hedin, M., 2016. Phylogenomic analyses resolve an ancient trichotomy at the base of Ischyropsalidoidea (Arachnida, Opiliones) despite high levels of gene tree conflict and unequal minority resolution frequencies. Mol. Phylogenet. Evol. 95, 171–182.

Rokas, A., Williams, B., King, N., Carroll, S., 2003. Genome-scale approaches to resolving incongruence in molecular phylogenies. Nature 425, 798–804.

Salichos, L., Rokas, A., 2013. Inferring ancient divergences requires genes with strong phylogenetic signals. Nature, 497, 3627–331.

Salichos, L., Stamatakis, A., Rokas, A., 2014. Novel information theory-based measures for quantifying incongruence among phylogenetic trees. Mol. Biol. Evol. 31, 1261–1271.

Sanderson, M.J., 1989. Confidence limits on phylogenies: the bootstrap revisited. Cladistics 5, 113–129.

Sayyari, E., Mirarab, S., 2016. Fast coalescent-based computation of local branch support from quartet frequencies. Mol. Biol. Evol. 33, 1654–1668.

Sayyari, E., Whitfield, J.B., Mirarab, S., 2017. Fragmentary gene sequences negatively impact gene tree and species tree reconstruction. Mol. Biol. Evol. 34, 3279–3291.

Shen, X., Hittinger, C.T., Rokas, A., 2017. Contentious relationships in phylogenomic studies can be driven by a handful of genes. Nature Ecol. Evol. 1, 0126.

Siddall, M., 1995. Another monophyly index: revisiting the jackknife. Cladistics 11, 33–56.

Simmons, M.P., 2017. Mutually exclusive phylogenomic inferences at the root of the angiosperms: *Amborella* is supported as sister and Observed Variability is biased. Cladistics 33, 488–512.

Simmons, M.P., Gatesy, J., 2015. Coalescence vs. concatenation: sophisticated analyses vs. first principles applied to rooting the angiosperms. Mol. Phylogenet. Evol. 91, 98–122.

Simmons, M.P., Sloan, D.B., Gatesy, J., 2016. The effects of subsampling gene trees on coalescent methods applied to ancient divergences. Mol. Phylogenet. Evol. 97, 76–89.

Simmons, M.P., Sloan, D.B., Springer, M.S., Gatesy, J., accepted. Gene-wise resampling outperforms site-wise resampling in phylogenetic coalescence analyses. Mol. Phylogenet. Evol.

Slowinski, J.B., Page, R.D.M., 1999. How should species phylogenies be inferred from sequence data? Syst. Biol. 48, 814–825.

Smith, S.A., Moore, M.J., Brown, J.W., Yang, Y., 2015. Analysis of phylogenomic datasets reveals conflict, concordance, and gene duplications with examples from animals and plants. BMC Evol. Biol. 15, 150.

Song, S., Liu L., Edwards, S.V., Wu, S., 2012. Resolving conflict in eutherian mammal phylogeny using phylogenomics and the multispecies coalescent model. Proc. Natl. Acad. Sci. USA 109, 14942–14947.

Springer, M.S., Gatesy, J., 2014. Land plant origins and coalescence confusion. Trends Plant Sci. 19, 267–269.

Springer, M.S., Gatesy, J., 2016. The gene tree delusion. Mol. Phylogenet. Evol. 94, 1–33.

Springer, M.S., Gatesy, J., 2018a. Pinniped diphyly and bat triphyly: more homology errors drive conflicts in the mammalian tree. J. Heredity 109, 297–307.

Springer, M.S., Gatesy, J., 2018b. Delimiting coalescence genes (c-genes) in phylogenomic datasets. Genes 9, 123.

Springer, M.S., Gatesy, J., 2018c. On the importance of homology in the age of genomics. Systematics and Biodiversity 16, 210–228.

Struck, T.H., Purschke, G., Halanych, K.M., 2006. Phylogeny of Eunicida (Annelida) and exploring data congruence using a partition addition bootstrap alteration (PABA) approach. Syst. Biol. 55, 1–20.

Vachaspati, P., Warnow, T., 2015. ASTRID: Accurate Species TRees from Internode Distances. BMC Genomics 16, S3.

Xi, Z., Liu, L., Davis, C.C., 2016. The Impact of missing data on species tree estimation. Mol. Biol. Evol. 33, 838–860.

Xi, Z., Liu, L., Rest, J.S., Davis, C.C., 2014. Coalescent versus concatenation methods and the placement of *Amborella* as sister to water lilies. Syst. Biol. 63, 919–932.

Zhang, C., Sayyari, E., Mirarab, S., 2017. ASTRAL-III: increased scalability and impacts of contracting low support branches. RECOMB International Workshop on Comparative Genomics, ed., J. Meidanis and L. Nakhleh. London, Springer: 53–75.

Zhong, B., Betancur-R, R., 2017. Expanded taxonomic sampling coupled with gene genealogy interrogation provides unambiguous resolution for the evolutionary root of angiosperms. Genome Biol. Evol. 9, 3154–3161.

Zhong, B., Liu, L., Yan, Z., Penny, D., 2013. Origin of land plants using the multispecies coalescent model. Trends Plant Sci. 18, 492–495

Zhong, B., Liu, L., Penny, D., 2014. The multispecies coalescent model and land plant origins: a reply to Springer and Gatesy. Trends Plant Sci. 19, 270–272.

Zwickl, D.J., Hillis, D.M., 2002. Increased taxon sampling greatly reduces phylogenetic error. Syst Biol. 51, 588–598.

